# A reproducibility-audit framework for generalizable versus dataset-specific molecular transition boundaries in Alzheimer’s disease

**DOI:** 10.64898/2026.08.24.746808

**Authors:** YoungOuk Kim, WooMyung Heo, Se Jin Park, YoungChul Kim, Ye Eun Cho

## Abstract

Molecular staging of Alzheimer’s disease (AD) increasingly defines transition boundaries along single-cell pseudo-progression trajectories, yet whether such boundaries reproduce across brain regions, cohorts and molecular modalities is rarely tested. We present a permutation-controlled audit that combines nine boundary-detection algorithms with a fixed marker panel and four orthogonal reproducibility axes—algorithmic consensus, region, cohort and modality. On synthetic data with planted ground-truth boundaries the audit reaches 100% sensitivity and 94% specificity, rejecting four distinct artefact classes each by a different axis. Applied to the Seattle Alzheimer’s Disease Brain Cell Atlas middle temporal gyrus, it localizes a transition that is robust across algorithms and recovered in most cell types but does not generalize: its leading marker is attenuated or absent in prefrontal cortex, entorhinal cortex and cerebrospinal fluid, and an apparent cross-region conservation of glial metabolic genes proves to be a global-expression offset rather than a shared program. The same audit nonetheless certifies an externally validated marker (astrocytic PTGDS) as reproducible across regions and modalities, showing that it separates generalizable anchors from dataset-specific ones rather than rejecting all signals. We provide this four-axis audit as a transferable, code-available standard to apply before a trajectory boundary is read as a biological stage, in AD and other progressive proteinopathies.

**Highlights:**

- A four-axis audit tests whether AD molecular-staging boundaries reproduce
- On synthetic ground truth it reaches 100% sensitivity and 94% specificity
- A robust MTG transition fails to generalize across regions, cohorts and CSF
- Apparent cross-region metabolic conservation is a global-expression artefact

*In brief:* Kim et al. develop a permutation-controlled, multi-algorithm audit that tests whether molecular transition boundaries in Alzheimer’s disease reproduce across regions, cohorts and modalities. Validated on synthetic ground truth, the audit separates a generalizable anchor (PTGDS) from a dataset-specific signal and a global-expression artefact, providing a reusable control for trajectory-based molecular staging.

*Motivation:* Single-cell atlases increasingly place disease stages at molecular transition boundaries along pseudo-progression trajectories, and the marker that anchors such a boundary is often read as biologically meaningful. Yet a boundary identified in one atlas need not recur in another region, cohort or molecular modality, and a large single-dataset effect can reflect global transcriptional amplitude rather than a specific program. No standard procedure exists to decide whether a trajectory boundary generalizes before it is interpreted. We therefore built a transferable, permutation-controlled audit with measurable operating characteristics that makes this decision explicit, and asked what it implies for one widely used atlas.

## Introduction

Defining biologically meaningful stages of Alzheimer’s disease (AD) is a central goal of contemporary neurodegeneration research, because disease mechanisms and therapeutic responses vary substantially across the disease continuum[1,2,3]. Biomarker-based frameworks already tie diagnosis and clinical-trial enrolment to staged definitions of the disease[1,3], so recovering the molecular events that demarcate one stage from the next is of more than descriptive interest. Single-nucleus and single-cell atlases now order individual nuclei along continuous pseudo-progression axes, and a rapidly growing literature uses these axes to define molecular “stages” or transition boundaries for biological interpretation and clinical stratification[4,5,6]. The appeal is intuitive: a transition boundary promises a compact, mechanistically interpretable coordinate at which a cell population changes regime, against which markers, genetic risk and candidate therapeutic windows can be aligned.

Three assumptions usually remain implicit: that a boundary found in one brain region reflects a brain-wide event; that a boundary found in one cohort generalizes to others; and that a marker reported as a boundary’s anchor is itself a reproducible disease signal rather than a feature of one dataset. None is trivial. Regional heterogeneity is a defining feature of AD neuropathology, with stereotyped but spatially staggered tau spread and region-specific vulnerability[7,8], and cross-cohort reproducibility of single-nucleus signatures is frequently limited by donor number, demographics and platform[9,10]. Single-cell differential-expression methods that treat individual nuclei as independent replicates are biased toward highly expressed genes and inflate false discoveries unless expression is aggregated to the donor level (pseudobulk)[11]. Boundary detection is itself methodologically fragile—segmented regression, clustering, change-point and trajectory methods frequently disagree on the same data[12,13,14]—so a consensus across methods evaluated against an empirical null is a more conservative basis for any claim than a single algorithm. These three assumptions are rarely tested together, yet each can fail on its own: a boundary may be genuine within a region but regionally restricted, reproducible in one cohort but not another, or robust as a boundary even when its nominal anchor marker is dataset-specific. Conflating them risks promoting an idiosyncratic feature of one dataset to the status of a brain-wide molecular stage.

Reactive astrocytes provide an informative molecular readout of this process. The canonical neurotoxic reactive-astrocyte program (GFAP, SERPINA3, complement, STAT3) is induced in AD[15,16,17], disease-associated astrocyte and microglial states have been catalogued[18,19], and the dominant mechanistic model places neurotoxic reactive astrocytes downstream of activated microglia[15,20]. Reactive astrocytes are not a single phenotype but a graded, context-dependent continuum spanning homeostatic and multiple reactive identities[16,17], and the disease-associated programs reported across studies overlap only partially[18,21]; whether any one program marks a defined position on the disease trajectory, rather than a generic stress response, is therefore an empirical question. Reactive astrocytes were chosen as a biologically informative test case because their disease-associated programs are widely reported across AD cohorts yet remain incompletely conserved across studies. We adopt this ordering and make no claim that astrocytes are an upstream causal origin of any transition reported here.

Here we develop a permutation-controlled, multi-algorithm consensus framework and use it to ask, deliberately conservatively, whether molecular transition boundaries and their proposed anchor markers remain reproducible across brain regions, cohorts and molecular modalities. Building on our earlier single-marker PTGDS–LCN2 boundary[22], we apply the framework to SEA-AD middle temporal gyrus and prefrontal cortex, an independent entorhinal cortex cohort and cerebrospinal-fluid proteomes, with explicit global-expression correction. We find that a transition can be robustly localized within one region yet shows substantial regional and modality dependence, that individual markers pass or fail external validation on their own evidence, and that distinguishing generalizable transitions from region-specific or artefactual ones requires the external and statistical controls this framework provides. The central claim is therefore not about any particular marker but about boundaries themselves: a molecular transition can be method-robust within a region yet region- and modality-specific rather than an intrinsic, brain-wide biological object. This distinction matters because transition boundaries are increasingly used to interpret disease mechanisms and define molecular stages from single-cell atlases[4,5,6]. Although consensus change-point detection and permutation testing are each established individually, combining them into a systematic, externally validated reproducibility audit of trajectory boundaries is rarely done, and to our knowledge has not previously been applied to the SEA-AD atlas at this scale. Our goal is not to reject boundaries, but to distinguish biologically reproducible boundaries from those whose apparent stability is dataset-dependent.

## Results

### Consensus framework localizes a reproducible MTG transition

We curated a 44-marker panel spanning A1/A2 reactive-astrocyte signatures, inflammation, vascular endothelium, AD risk genes, iron/oxidative stress, neurotrophic factors and core AD proteins (Table S1), fixed a priori and held constant across all cohorts; PTGDS, LCN2 and MAPT were deliberately excluded to permit independent boundary detection. Analysing 67,419 MTG astrocyte nuclei ordered along the continuous pseudo-progression score (CPS ≥ 0.1) and aggregated into nine CPS bins, we applied nine algorithms spanning four mathematical families—hierarchical clustering, partitioning, piecewise/segmented regression and change-point detection (19 algorithm–component combinations)—to the leading principal components (PC1– PC3) of the bin-aggregated marker matrix, combining their calls into a consensus localization. The algorithms localized a primary transition at CPS 0.207, recovered concordantly by Ward.D2 (variance-minimizing agglomerative) hierarchical clustering (k = 3), k-means, variance-jump and trajectory-inflection methods. A second, weaker candidate boundary fell near CPS 0.39–0.42 (dynamic-programming change-point on PC2 at CPS 0.388), coinciding with the ADNI (Alzheimer’s Disease Neuroimaging Initiative) Mini-Mental State Examination (MMSE)-26 transition and the clinical mild-cognitive-impairment (MCI) threshold. Across 1,000 random 44- gene panels, the PC1 piecewise breakpoint spanned CPS 0.26–0.75 (median 0.45) and never fell within ±0.05 CPS of 0.21 (0/1000), establishing the primary transition as exceptionally panel-specific. The change-point methods instead localized a mid-trajectory split near CPS 0.54, but this position was reproduced by 40% of random panels and is therefore non-specific, whereas the weaker secondary near CPS 0.39–0.42 showed only intermediate specificity (empirical P = 0.127); the full per-algorithm boundary distribution is provided in Table S2. Separating a panel-specific transition from these non-specific or weakly specific splits is precisely the dataset-dependent structure the framework is designed to resolve (Figure 1).

**Figure 1.**
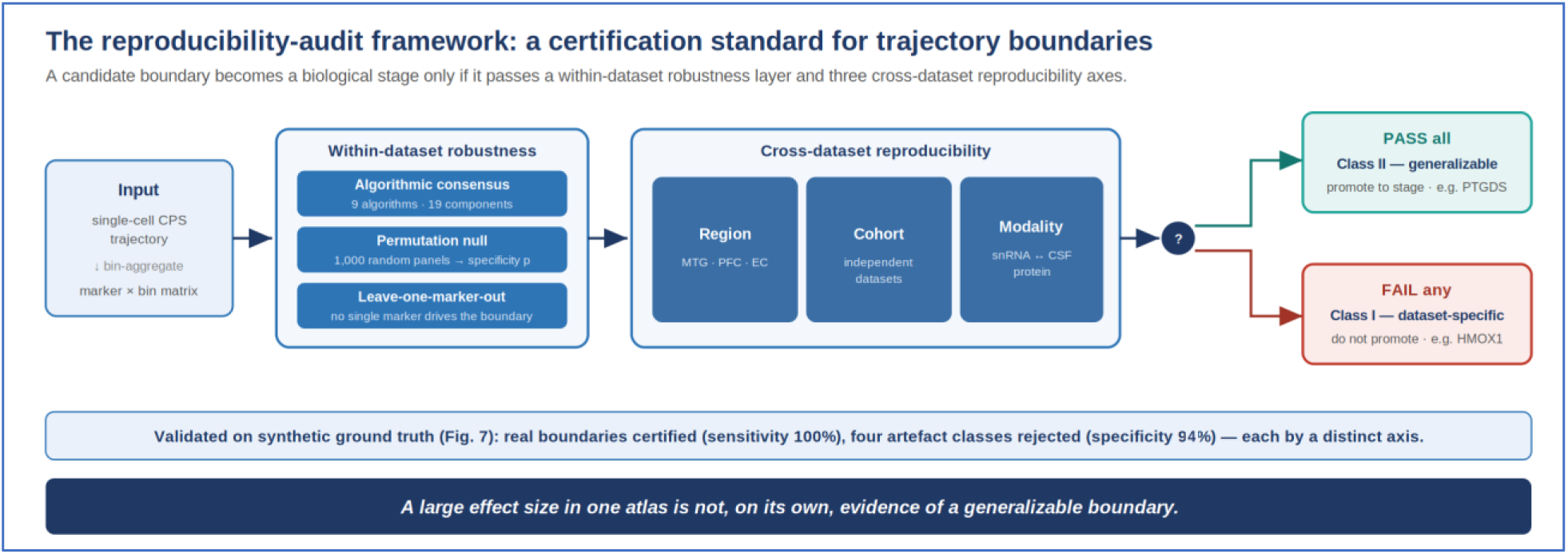
The reproducibility-audit framework. A candidate molecular transition boundary, detected on a single-cell pseudo-progression (CPS) trajectory and aggregated into a marker × bin matrix, is first held to a within-dataset robustness layer—multi-algorithm consensus (nine algorithms, 19 components), a random-panel permutation null that scores panel-specificity, and leave-one-marker-out stability—and then to three cross-dataset reproducibility axes: region, cohort and molecular modality. The region axis tests a pan-regional (brain-wide) claim rather than requiring identical recurrence across regions: because regional heterogeneity is expected in Alzheimer’s disease, a region-specific boundary is read as regional rather than universal, and it is an over-broad universality claim—not the boundary itself—that the axis refutes. Boundaries passing every test are promoted as generalizable Class II anchors (e.g. astrocytic PTGDS); those failing any test are dataset-specific Class I signals or global-expression artefacts (e.g. HMOX1). The framework is validated on synthetic data with planted ground-truth boundaries, where it certifies genuine boundaries (sensitivity 100%) and rejects four artefact classes (specificity 94%), each caught primarily by a distinct axis (Figure 7). A large effect size in a single atlas is not, on its own, evidence of a generalizable boundary.

The transition is defined by coordinated reactive-astrocyte activation: at CPS 0.21, A1 markers rose together (the boundary effect Δz, i.e. the change in mean z-score across the transition: GFAP +0.48, FKBP5 +0.47, BCL2 +0.35, SERPINA3 +0.20, STAT3 +0.17). The single largest change in the dataset was a decrement in HMOX1 (heme oxygenase-1, an iron/oxidative-stress response gene; Δz = −0.60)[23,24], which we evaluate against external data below. Leave-one-marker-out analysis showed that the CPS-0.21 boundary is recovered unchanged when HMOX1 is removed—it is co-defined by GFAP, FKBP5 and BCL2—so the transition does not depend structurally on any single gene (Figure 2).

**Figure 2.**
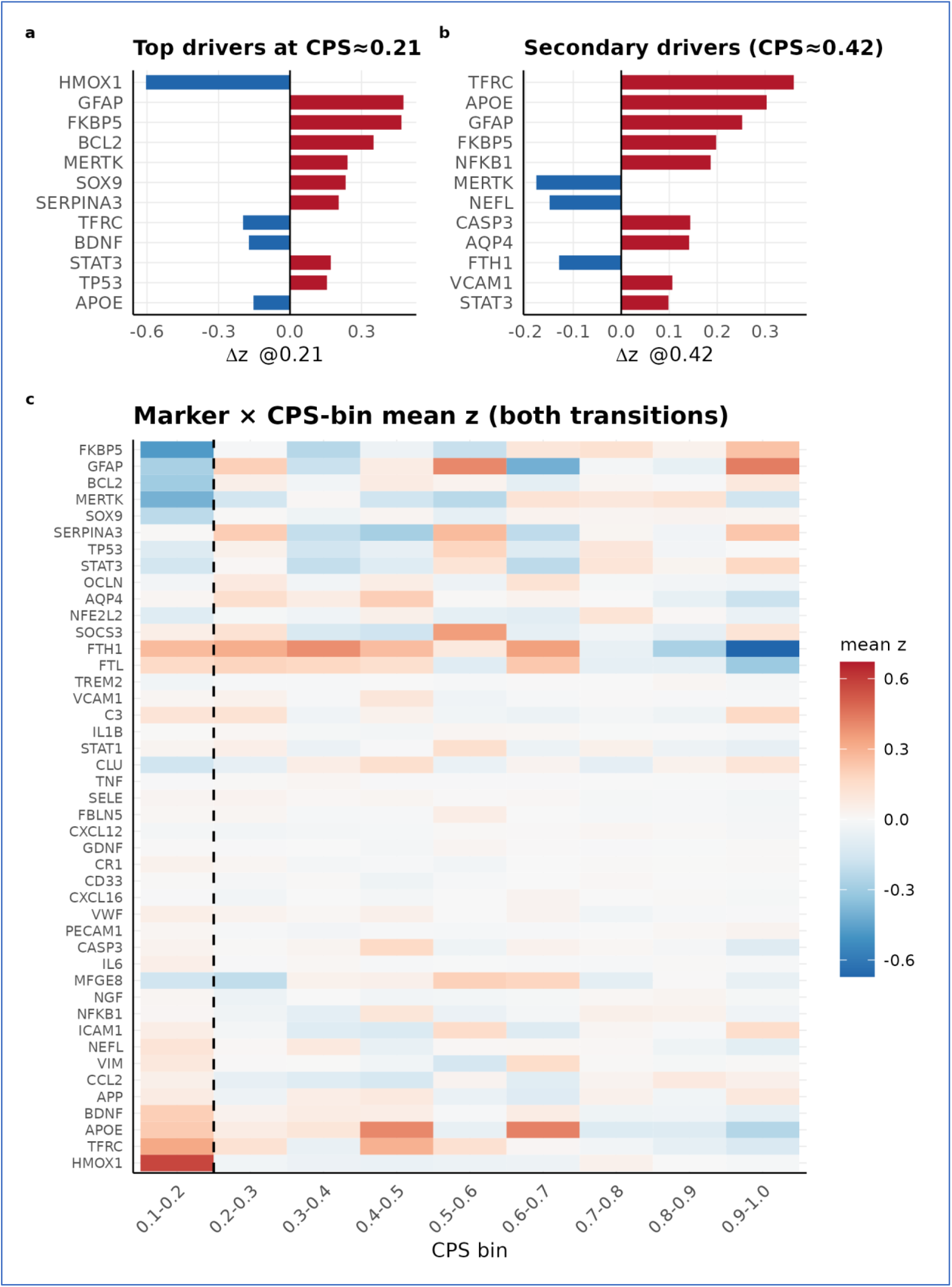
A multi-algorithm consensus localizes a reproducible reactive-astrocyte transition in SEA- AD MTG. **(a)** Top drivers of the CPS≈0.21 transition (GFAP/FKBP5/BCL2 up; HMOX1 the largest single decrement, shown but treated as a candidate for external testing). **(b)** Drivers of the secondary MCI-stage transition at CPS≈0.42. **(c)** Marker × CPS-bin mean-z heatmap across the pseudo- progression continuum, showing both transitions; the CPS-0.21 boundary was recovered by 0/1000 expression-matched random panels within ±0.05 CPS and is unchanged on leave-one-marker-out removal of HMOX1 (co-defined by GFAP/FKBP5/BCL2).

### The transition is recovered across MTG cell populations

Applying the identical panel and framework to seven further MTG cell populations, the CPS-0.21 transition was recovered in seven of eight populations (5–9 algorithms each: astrocytes, excitatory and inhibitory neurons, endothelial cells, oligodendrocyte precursor cells (OPCs), microglia and vascular leptomeningeal cells (VLMCs)), with oligodendrocytes the sole outlier (Figure S1). FKBP5 upregulation was the most broadly shared driver—strongest in microglia (Δz +1.06), OPCs (+0.65) and astrocytes (+0.47) but essentially absent in inhibitory neurons (|Δz| < 0.07)[25,26]—whereas the HMOX1 decrement (astrocyte Δz −0.60) and GFAP induction (+0.48) remained astrocyte-specific (Table S3). We describe this as a coordinated, cross-cell-type event within MTG—a within-region property distinct from the brain-wide generalization tested below. Inherited AD risk aligned with microglia rather than astrocytes at this boundary: a pre-registered set of 41 detected AD genome-wide association study (GWAS) risk genes was coordinately upregulated in microglia (mean |Δz| = 0.088 versus expression-matched null 0.070; empirical P = 0.029, nominal and ≈ 0.06 after correction for the two cell types tested) but showed no convergence in astrocytes (P = 0.22), consistent with the predominantly microglial genetic architecture of AD[27,28] and positioning the astrocyte signal as a downstream, post-microglial reactive readout rather than a genetic driver (Figure S2).

### Cross-region and modality testing show limited conservation

To test whether the CPS-0.21 transition is brain-wide, we re-ran the identical pipeline in SEA-AD dorsolateral prefrontal cortex (Brodmann area 9): the transition was markedly attenuated (one of eight cell types), and the HMOX1 change fell from Δz −0.60 (MTG) to ≈ −0.06 (A9). In an independent entorhinal cortex (EC) cohort[29,30] (Leng et al. 2021; GSE147528; 10 male ApoE3/3 donors) processed with an emptyDrops + Harmony + edgeR pseudobulk pipeline, high-expression markers moved opposite to, or independently of, the MTG signature: FTH1 increased pan-cellularly (astrocytes ΔlogFC +0.54, P = 7×10⁻⁴), opposite to the MTG iron-storage decline[31,32], while TFRC and GFAP were flat. A within-cohort positive control (GFAP-high versus GFAP-low astrocytes; GFAP ΔlogFC +12.5) recovered downregulation of all eight canonical homeostatic genes, six significantly (NRXN1 −0.67, SLC1A2 −0.64, PTN −0.52, GPC5 −0.51, CADM2 −0.40, SLC1A3 −0.35; false discovery rate (FDR) < 0.05), confirming that the pipeline correctly recovers reactive de-homeostasis; critically, because the pipeline detects a coordinated reactive-astrocyte signal in these same ten donors, the non-recovery— indeed reversal—of the MTG panel is an informative negative rather than an artefact of limited power, and region- and cohort-specificity is therefore a reproducible biological observation, not a technical failure (Figure 3; Table S4).

**Figure 3.**
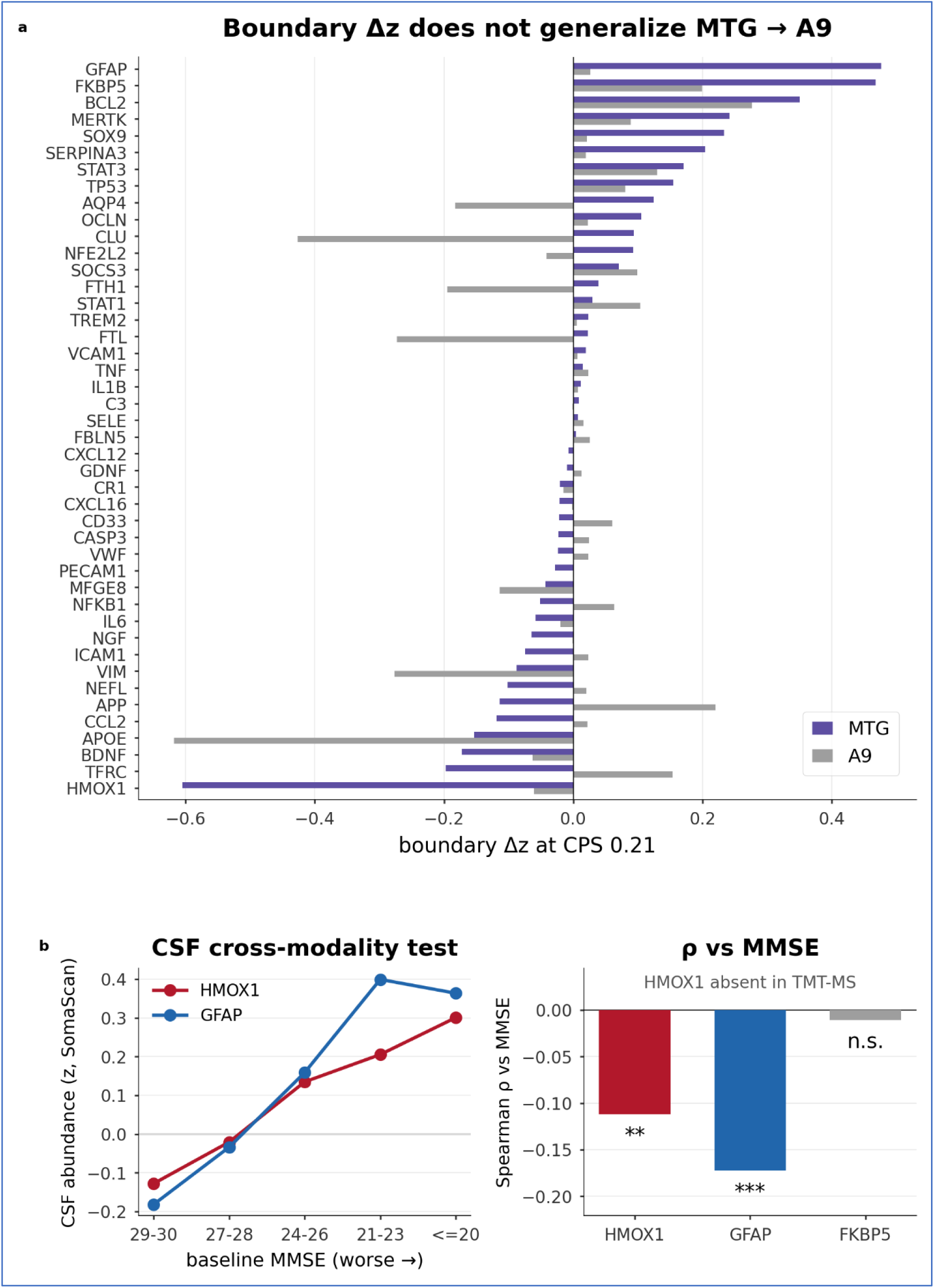
The transition and its lead marker do not generalize. **(a)** Panel-wide boundary Δz in MTG vs A9 astrocytes: the MTG reactive signature (FKBP5/GFAP/BCL2 up; HMOX1 −0.60 down) is strongly attenuated in A9 (HMOX1 −0.06). **(b)** CSF cross-modality test of HMOX1 vs GFAP: HMOX1 is undetected in TMT-MS and weakly reversed in SomaScan (ρ=−0.11), whereas GFAP rises consistently with impairment (ρ=−0.17). In an independent EC cohort the panel runs opposite to MTG (pan-cellular FTH1 increase, ΔlogFC +0.54; flat TFRC/GFAP), with an EC positive control (8/8 canonical homeostatic genes down in GFAP-high astrocytes) confirming pipeline validity.

This regional difference reflects astrocyte composition rather than a within-subtype reversal. Comparing the six SEA-AD astrocyte supertypes between regions at the donor level (centred-log-ratio proportions, Wilcoxon rank-sum, n = 84 donors), composition differed markedly: the MTG-enriched Astro_1 supertype was ∼3.8-fold more abundant in MTG than A9 (median 7.2% versus 1.9%, P_adj = 8×10⁻¹⁶), whereas the homeostatic-dominant Astro_2 was relatively enriched in A9 (64.7% versus 70.5%, P_adj = 2.5×10⁻³). The attenuation of the MTG marker boundary in A9 therefore coincides with depletion of the MTG-enriched Astro_1 supertype rather than a sign reversal within a shared astrocyte population, consistent with a fixed panel sampling different subtype mixtures across regions.

The lead MTG marker behaved similarly under direct external testing. In cerebrospinal fluid (CSF), HMOX1 was undetected (ADNI Emory tandem mass tag mass spectrometry, TMT-MS) or, where measured (ADNI SomaScan 7K; n = 724), showed only a weak association with cognition in the direction opposite to the brain signal (Spearman ρ = −0.11, P = 3.5×10⁻³), whereas the reactive marker GFAP behaved consistently and more strongly (ρ = −0.17, P = 4×10⁻⁶; higher CSF GFAP with greater impairment) and FKBP5 was flat (ρ = −0.015). HMOX1 thus showed limited reproducibility across regions and modalities, and we re-anchor the biological description of the transition on GFAP (Figure 3; Table S5). By contrast, a companion- study marker (astrocytic PTGDS) passed the same region- and modality-tests—conserved across human, zebrafish and murine systems and reproduced in external bulk-tissue proteomic cohorts[22,33]—providing a positive control that the framework accepts genuinely generalizable signals rather than rejecting all candidates (Table S6).

### Apparent cross-region metabolic conservation is a global-expression artefact

Energy-metabolism failure is mechanistically implicated in AD[34,35,36], so we asked whether a glial lactate/glycolysis supply program shows a conserved transition across regions, analysing 21 metabolic-supply genes per cell type in MTG and A9. An uncorrected analysis appeared to support conservation (eight genes sharing the same sign of change), but this was an artefact of global transcriptional activity: at the MCI boundary, A9 showed a near-uniform negative shift across all cell types (up to 100% of genes negative in astrocytes). After per-cell-type global- expression correction—subtracting, within each region and cell type, the mean effect across all genes, a standard control for the differences in global transcriptional activity, total counts and detection rate that otherwise inflate cross-gene and cross-region correlations—only eight of the 21 genes shared the same sign of change (13 divergent), leaving no coherent conserved metabolic transition (Figure 4). The procedure is applied identically to every gene and is symmetric by construction: it removes only the offset shared across all genes within a cell type and would leave a genuinely gene-specific conserved signal intact, so it functions as a confound control rather than as a device for removing inconvenient signal. A genuine global decline in transcriptional output during disease progression may itself be real biology, but a claim that a specific glial metabolic-supply programme is conserved across regions requires that programme to rise above this shared trend, which it does not once the offset is removed. As an empirical negative control, a panel of 14 constitutive housekeeping genes remained essentially flat along the trajectory both before and after correction (mean |Δz| ≈ 0.055 in both MTG and A9, unchanged by correction; Figure S3), confirming that the procedure removes only the shared global offset and leaves gene-specific signal intact. A naive cross-region comparison of module-average effects can therefore manufacture apparent conservation, and global-expression correction should be a routine control. Crucially, this artefact concerns only naive cross-region module averaging and does not negate genuine within-region metabolic programs—on the contrary, it underscores why the within-region, donor-level designs of the companion studies are necessary. Those analyses resolve a real astrocytic lactate-export decline (MCT4/SLC16A3)[37] with an internal dissociation control (export genes fall while V-ATPase is preserved), while neuronal MCT2 (SLC16A7) import is preserved—indicating the bottleneck is astrocytic export rather than neuronal import; the present framework therefore corroborates, rather than contradicts, those within-region findings.

**Figure 4.**
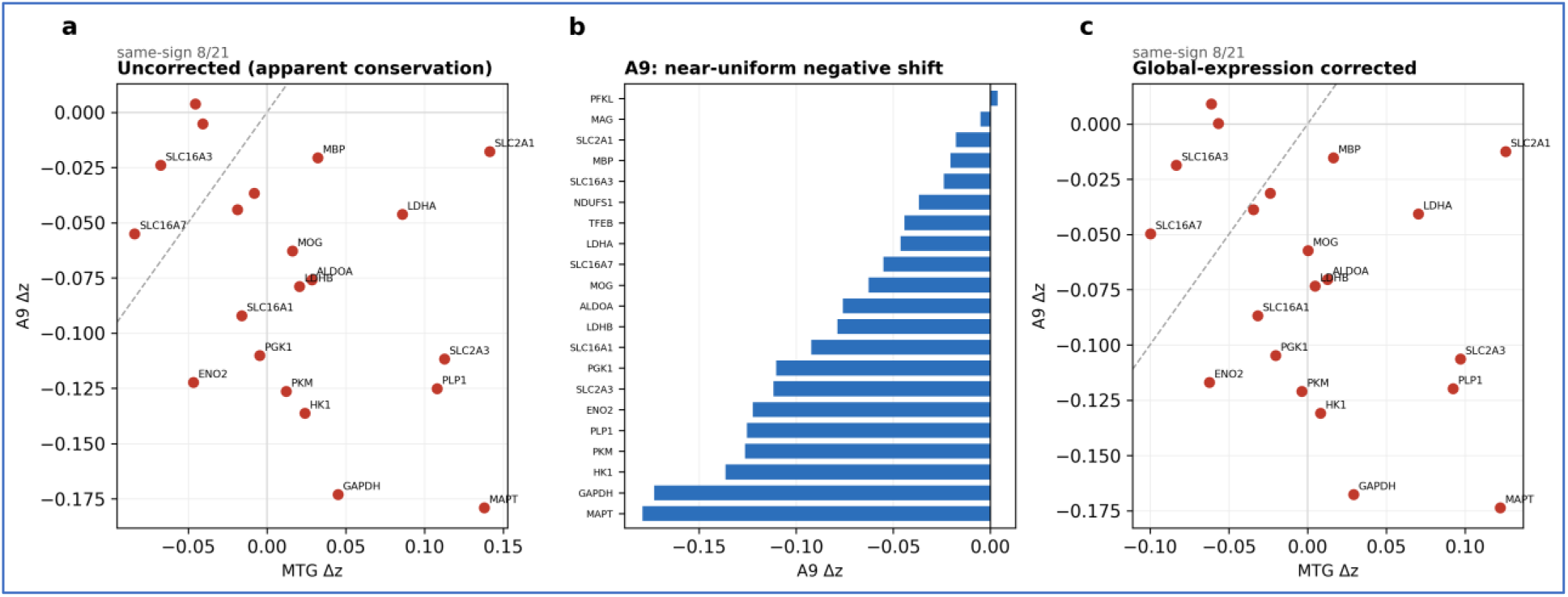
Apparent metabolic conservation is a global-expression artefact of cross-region module averaging. **(a)** Uncorrected metabolic-gene module averages appear similar between MTG and A9. **(b)** A9 shows a near-uniform negative shift across all cell types. **(c)** After global-expression correction, only 8 of 21 genes remain same-sign across MTG and A9 (13 divergent). A within-region, donor-level dissociation analysis (companion study) resolves a genuine astrocytic lactate-export decline (MCT4) against preserved V-ATPase, while neuronal MCT2 import is preserved.

### Partial discordance between brain and CSF trajectories

Applying the framework to CSF proteomes (ADNI SomaScan, n = 343; ADNI Emory TMT-MS, n = 1,104 of 1,105), algorithm outputs again clustered into two regions, with a clinically aligned boundary near MMSE 26. However, brain (single-nucleus RNA, snRNA) and CSF (protein) trajectories were not uniformly concordant at the marker level—HMOX1 in particular showed a sign reversal between compartments—and several intracellular drivers (HMOX1, FKBP5, STAT3) fell below CSF detection in TMT-MS. Because the MMSE-26 boundary is anchored to a clinical threshold by construction, we report the brain–CSF discordance as a finding in its own right—a caution for cross-modality biomarker translation—rather than as multi-platform confirmation of a single boundary (Figure 5a).

**Figure 5.**
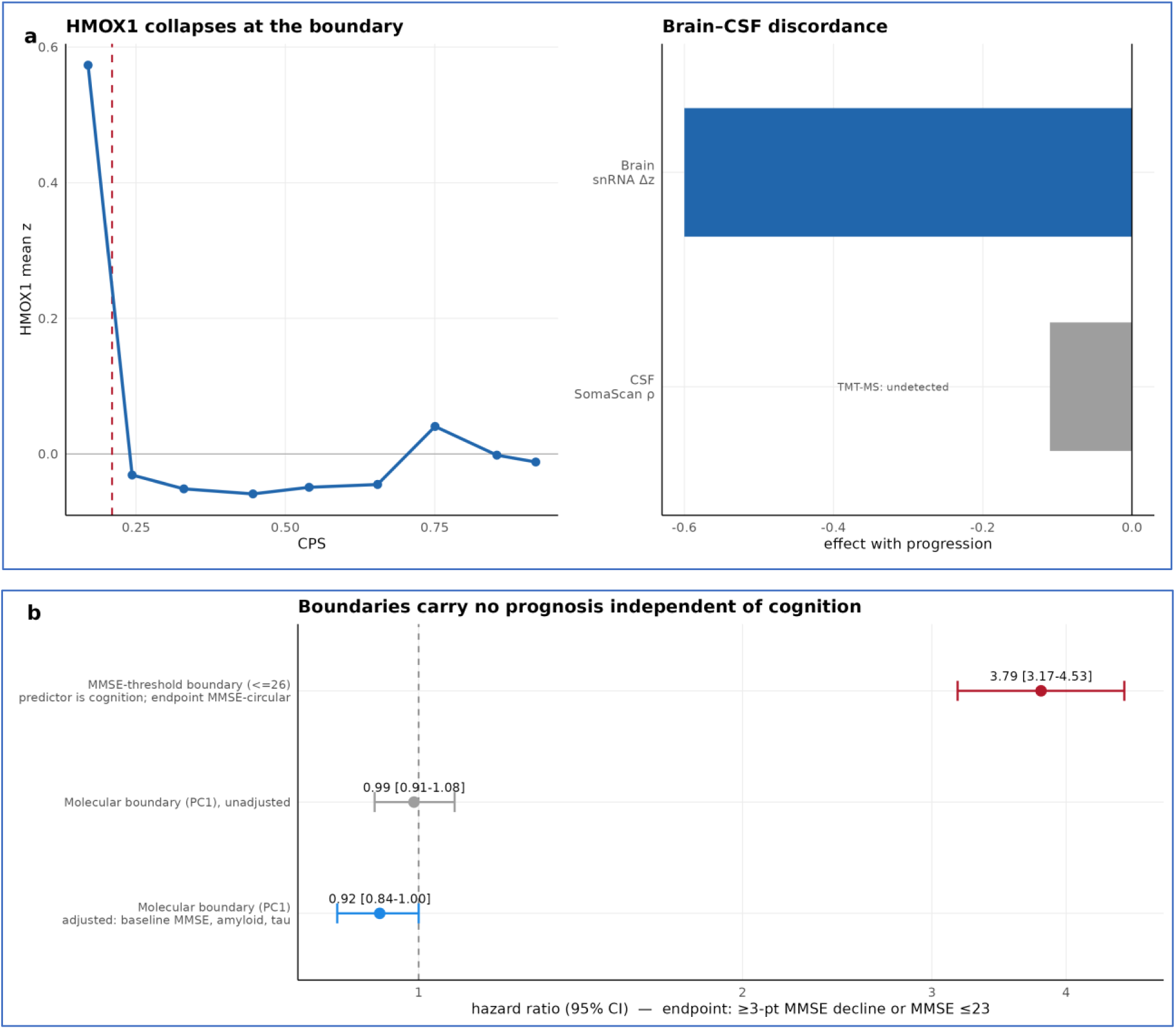
Brain–CSF discordance and absence of cognition-independent prognosis. **(a)** HMOX1 changes in opposite directions in brain versus CSF. **(b)** A cognition-threshold survival model is circular (MMSE ≤ 26 HR 3.79, 95% CI 3.17–4.53), whereas the molecular panel score is essentially null after adjustment for baseline cognition and amyloid–tau status (HR 0.92 per SD, 95% CI 0.84–1.00; 0.99 unadjusted).

### Boundaries add limited prognostic value beyond cognition

We examined prognostic value in the Emory cohort (n = 1,105 with matched baseline cognition, CSF amyloid–tau and panel proteomics; 527 events; endpoint ≥ 3-point MMSE decline or MMSE ≤ 23). A boundary operationalised by a baseline MMSE threshold (≤ 26) returned a large hazard ratio (HR = 3.79, 95% CI 3.17–4.53), but because this predictor is a cognitive threshold and the endpoint is itself MMSE-based, the association is largely circular. Defined molecularly instead (the first principal component of the available CSF panel proteins), the boundary added essentially no prognostic value beyond baseline cognition and amyloid–tau status (HR 0.92 per s.d., 95% CI 0.84–1.00, P = 0.061 adjusted; HR 0.99, P = 0.78 unadjusted). That a cross-sectional boundary carries no prognostic information independent of cognition is itself an outcome of the audit—the framework correctly flagging a claim that does not generalize to longitudinal decline—rather than a shortcoming of it. The MMSE-aligned boundary is therefore best understood as a molecular recovery of an already-established clinical threshold; this concerns prognosis (rate of decline) rather than staging (position on the trajectory), and the boundaries are cross-sectional staging constructs rather than independent prognostic markers (Figure 5b).

### Robustness and donor-level (pseudobulk) validation controls

Within MTG, the boundaries were stable to analytical choices: a bin-shift analysis (±0.08 CPS), alternative MMSE binning, leave-one-marker-out (28 iterations), and APOE ε4 / sex stratification[38] preserved the two-boundary architecture, and two assumption-light methods (variance-jump; Ward clustering) independently localised the CPS-0.21 transition. Re- introducing PTGDS and LCN2 in a 39-marker panel[39] left the primary boundary essentially unchanged (CPS 0.207) while the weaker secondary boundary was more labile (shifting within the CPS 0.29–0.42 range with panel composition), consistent with its lower permutation specificity; the previously reported PTGDS biphasic trajectory (vertex CPS ≈ 0.47) lies just beyond this secondary boundary, at the close of the compensatory phase; the multi-marker boundary and the single-marker PTGDS inflection thus localise to the same compensatory-to- vulnerable transition, so the present framework extends rather than contradicts the single-marker PTGDS finding (Figure 6).

**Figure 6.**
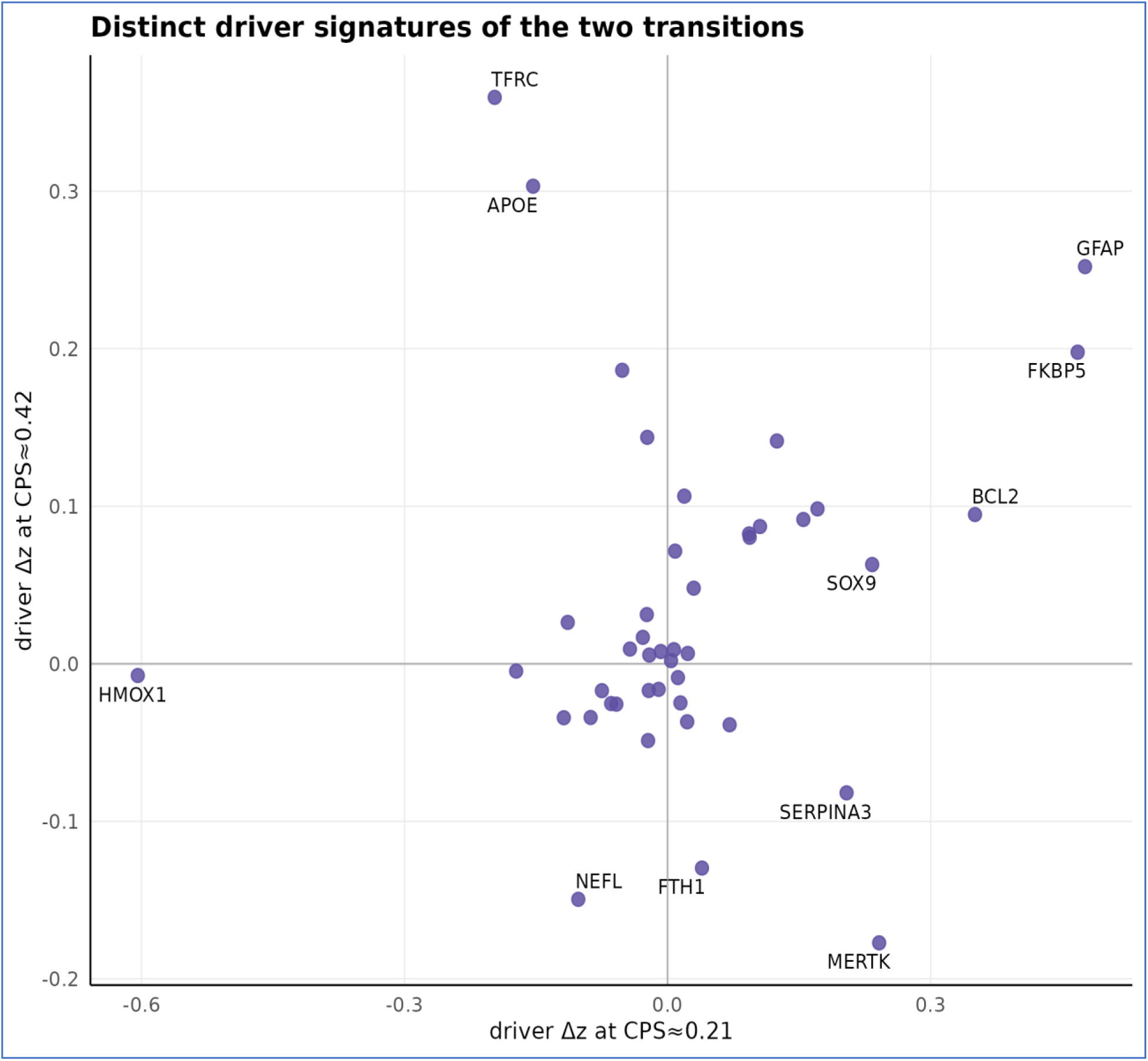
Distinct driver signatures of the two within-MTG transitions. Each point is a panel marker positioned by its boundary effect (Δz) at the primary transition (CPS≈0.21, x-axis) versus the secondary transition (CPS≈0.42, y-axis). The two transitions are driven by largely distinct marker sets: reactive- astrocyte markers (GFAP, FKBP5, BCL2) and the HMOX1 decrement separate along the primary axis, whereas a partly distinct set (e.g., TFRC, APOE) loads on the secondary axis.

Finally, we confirmed that donor-level (pseudobulk) testing is necessary rather than optional. Across 24 SEA-AD MTG cell types over a fixed 4,000-gene random universe, cell-level testing that treats individual nuclei as replicates inflated the number of differentially expressed genes (DEGs; FDR < 0.05) roughly four-fold relative to donor-level pseudobulk (edgeR; median 4.0×, up to ≈85× in cell types with few donors—for example 846 versus 10 genes), the inflation scaled with nucleus number (Pearson r = 0.52), and cell-level calls were enriched for highly expressed transcripts (18 of 24 cell types)—the documented signature of pseudoreplication. All cohort contrasts in this study therefore use donor-level pseudobulk, so that the region- and cohort- specificity reported above reflects biology rather than a replicate-counting artefact (Figure S4).

### A synthetic benchmark establishes the audit’s operating characteristics

The audit combines established components—consensus change-point detection, permutation testing and leave-one-marker-out robustness—so its value rests not on any single step but on whether the assembled gates reliably separate reproducible boundaries from artefacts. To measure this directly, we benchmarked the identical engine and gates on synthetic data in which the presence and location of a boundary are fixed by construction (Methods). Across five ground-truth regimes, the audit certified every planted panel-specific boundary (sensitivity 100%) and rejected all four artefact classes (specificity 94%), with each artefact excluded primarily by a different axis (Figure 7a). A global-expression offset—the synthetic analogue of the cross-region metabolic artefact above, in which every gene carries the same shift—passed the consensus- support gate but was reproduced by random panels and so failed the specificity test (median specificity P = 0.21 versus 0.01 for genuine panel-specific boundaries); null and monotonic-ramp panels failed for lack of consensus support; and a single-marker decoy that passed both support and specificity was caught only by leave-one-marker-out, where removing its one driver shifted the boundary by a median of 0.30 CPS against 0.00 for genuine boundaries (Figure 7b). The benchmark also shows why no single algorithm suffices: individual-method localization error ranged from 0.00 to 0.60 CPS, whereas the consensus localized the planted boundary to within 0.025 CPS and stayed within 0.022–0.025 CPS when any one of the nine algorithm families was withheld—so the result depends on agreement across methods, not on the specific algorithm set (Figure 7c,d). These operating characteristics are a property of the audit itself, independent of any biological interpretation, and provide a quantitative basis for the certifications reported above.

**Figure 7.**
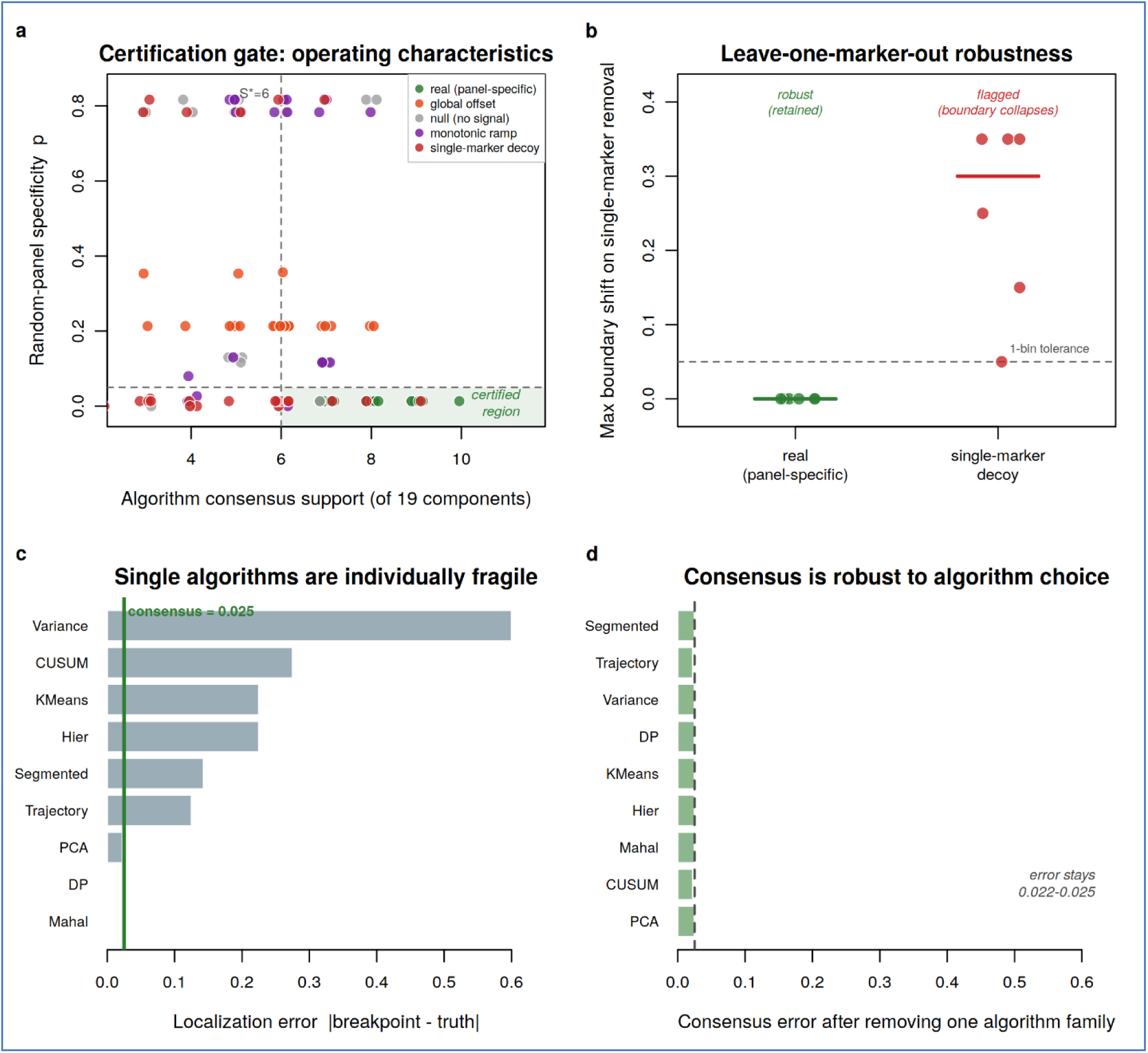
Synthetic benchmark with planted ground-truth boundaries. The identical detection engine and audit gates were applied to synthetic marker × bin matrices in which the presence and location of a boundary are known by construction (Methods). **(a)** Operating characteristics in the support × specificity plane: real panel-specific boundaries (green) fall in the certified region (consensus support ≥ S*, specificity P ≤ 0.05), whereas global-expression offsets, null and monotonic-ramp panels are excluded; a single-marker decoy intrudes into the region and is resolved in (b). **(b)** Leave-one-marker-out robustness: removing the single most influential marker leaves genuine boundaries unmoved (maximum shift 0.00 CPS) but collapses the decoy (median 0.30 CPS), beyond the one-bin tolerance. **(c)** Single-algorithm localization error against the planted boundary spans 0.00–0.60 CPS, whereas the multi-algorithm consensus (green line) localizes it to 0.025 CPS. **(d)** Leave-one-algorithm-out: the consensus error stays 0.022–0.025 CPS when any one of the nine algorithm families is withheld, so localization is not contingent on the exact algorithm set. Sensitivity 100%, specificity 94%, with each artefact class rejected primarily by a distinct axis.

## Discussion

Using a permutation-controlled, nine-algorithm consensus framework, we identified a reproducible reactive-astrocyte transition in the SEA-AD MTG, recovered across seven of eight cell types, and the identical audit applied to an independent microglia-specific marker panel localized a concordant transition (CPS 0.20; Mahalanobis, Ward, k-means and trajectory- inflection), confirming the framework is panel- and cell-type-agnostic (Figure S5). At this boundary, inherited AD genetic risk showed a nominal trend toward microglia rather than astrocytes—a pre-registered set of 41 AD-GWAS risk genes was coordinately upregulated in microglia (empirical P = 0.029, nominal; ≈ 0.06 after correction for the two cell types tested) but not in astrocytes—positioning the astrocytic transition as a downstream reactive readout rather than a genetic driver. Rather than a brain-wide molecular switch, however, the transition shows substantial regional and modality dependence: it is strongly attenuated in prefrontal cortex and is not reproduced in an independent entorhinal cortex cohort, where the same panel moves in the opposite direction. The principal contribution of this work is therefore not a single molecular boundary, nor any individual statistical technique, but their integration into a practical reproducibility benchmark for trajectory-based molecular staging: an empirically grounded, statistically controlled framework that separates molecular transitions which generalize from those that are region-specific or artefactual. Applying this standard, astrocytic PTGDS is adjudicated as a Class II anchor consistent with all four axes—an external positive control whose reproducibility is established in the companion study and external cohorts rather than discovered in the present panel; adjudicating externally proposed candidates is the framework’s intended function rather than a limitation, and an independent within-dataset signal—a nominal AD-GWAS trend in microglia (≈ 0.06 after correction)—provides suggestive supporting biology arising here.

As a concrete worked example, we ran the audit at the donor level in SEA-AD MTG on a panel of fourteen widely cited candidate markers, asking whether each reproduces its reported direction as a continuous transcriptional trajectory along CPS (Table S7). Only three—NPTX2, VGF and SCG2—did so (Spearman against CPS, FDR-significant, monotonic and stable across bin choices); the remaining eleven did not. The cerebrospinal-fluid prognostic markers NPTX1 and NPTXR[40] illustrate why: NPTX1 neuronal mRNA rises with CPS (ρ = +0.44) even as its CSF and brain protein fall with disease[40], a direct inversion between compartments, while NPTXR mRNA falls—neither is the flat, brain-preserved signal a protein-level reading would predict. The canonical glial markers GFAP, CHI3L1 and sTREM2/TREM2 showed no significant donor-level transcriptional trend despite robust protein-level reactivity. These divergences are not pipeline artefacts—NPTX2, the best-established member, reproduces cleanly (ρ = −0.58)[41,42]—but a consequence of the axis being interrogated: a marker certified in cerebrospinal-fluid protein need not, and frequently does not, reproduce as a brain transcriptional transition, and its direction can invert between compartments. The audit is bundled for single-cell application in the accompanying package, so that newly proposed markers can be classified— and, where they fail to generalize, flagged—as evidence accrues.

Two methodological results carry relevance beyond this dataset. First, the framework discriminates between two classes of candidate anchor marker by holding each to the same four orthogonal reproducibility axes—algorithmic consensus, region, cohort and molecular modality. Here the region axis is a falsification test of a pan-regional claim, not a requirement that a boundary recur identically across regions: regional heterogeneity is expected in Alzheimer’s disease, so a region-specific boundary is read as regional rather than artefactual, and a single well-powered counterexample suffices to refute an over-broad brain-wide claim. Class I markers anchor a boundary in one dataset but do not survive these tests: HMOX1, the single largest MTG change, is attenuated in prefrontal cortex, inverted or independent in entorhinal cortex, and undetected or sign-reversed in cerebrospinal fluid. Class II markers pass them: astrocytic PTGDS is conserved across species and external bulk-tissue cohorts in a companion study. That HMOX1 produces the most dramatic single-dataset change yet falls into Class I is the central illustration of the framework’s value—a large effect size in one atlas is not, on its own, evidence of a generalizable boundary, and external and statistical controls rather than effect magnitude decide the question (Table S6). Second, and most consequential, an apparent cross-region conservation of a glial metabolic-supply program dissolved after global-expression correction: prefrontal cortex showed a near-uniform negative shift across all cell types, and once this global activity was removed, conserved and divergent signals were balanced. A naive comparison of module-average effects across regions can therefore manufacture apparent conservation, and global- expression correction—like donor-level (pseudobulk) aggregation, which we confirm is required to avoid highly-expressed-gene bias and false-discovery inflation—should be a routine control in single-cell disease staging.

As a practical matter, the controls that separate generalizable from artefactual transitions can be assembled into a transferable protocol (Box 1). Four elements proved decisive here and are inexpensive to apply: a multi-algorithm consensus evaluated against a permutation null, which prevents any single change-point or trajectory-inference method[12,13,14] from dictating a boundary—extending resampling-based consensus clustering[43] from class discovery to boundary localization; donor-level (pseudobulk) aggregation, without which cell-level testing inflated differential-expression counts several-fold and over-weighted highly expressed genes[9,10,11]; per-cell-type global-expression correction, which dissolved an apparent metabolic conservation that had survived naive module averaging; and direct external testing of each candidate anchor in an independent region, cohort and molecular modality. None of these is novel in isolation, but applied together they convert a single-region observation into a claim whose scope is explicitly bounded—and, where a marker passes all four, into a more credible generalizable signal. We suggest this combination as a minimal standard for boundary-based molecular staging, in AD and in other progressive proteinopathies where trajectory boundaries are increasingly used for stratification. To our knowledge, such a systematic, permutation-controlled reproducibility audit of trajectory boundaries has not previously been reported for the SEA-AD atlas, and the protocol is portable to any pseudo-progression dataset. Unlike an ad-hoc pipeline, the assembled audit has measurable operating characteristics: on synthetic data with planted ground-truth boundaries it certified genuine boundaries and rejected four artefact classes—each caught by a different axis—at a sensitivity of 100% and a specificity of 94% (Figure 7), confirming that the axes are non-redundant rather than merely stacked.

### Box 1 A transferable reproducibility-audit protocol for trajectory boundaries

1. Localize the boundary by multi-algorithm consensus (≥9 algorithms spanning clustering, partitioning, segmented-regression and change-point families) and retain only boundaries that beat a permutation null (panel-specific; e.g., recovered by 0/1,000 expression-matched random panels).

2. Aggregate to donor level (pseudobulk) before differential testing, to avoid false-discovery inflation and highly-expressed-gene bias.

3. Apply per-cell-type global-expression correction before any cross-region conservation claim, removing the global transcriptional-amplitude offset that can manufacture apparent conservation.

4. Externally test every candidate anchor across four orthogonal axes: algorithmic consensus, region, cohort and molecular modality.

Verdict: anchors passing all four axes are generalizable (Class II); those failing any axis are dataset-specific (Class I). See Table S6 for the worked HMOX1 (Class I) versus PTGDS (Class II) comparison.

Biologically, the regional staggering we observe—sharp in MTG, attenuated in prefrontal cortex, inverted in entorhinal cortex—is consistent with independent multi-region evidence that astrocytic glycolytic compensation peaks at different disease stages across regions (early in entorhinal, intermediate in temporal, late in prefrontal cortex)[4,44]. Region-specificity is thus not mere non-replication but a biologically interpretable consequence of regionally staggered trajectories that a single-region boundary projected brain-wide would misrepresent; this scoping concerns marker- and module-average effects and is not in conflict with the broad cross-region similarity in coarse cell-composition dynamics reported by recent atlases. Importantly, delimiting where single-dataset signals generalize refines—rather than retracts—the within-region programs established by stricter designs, including the companion astrocytic lactate-export decline and the conserved staging marker. The brain–CSF discordance is likewise reported as a finding in its own right—a caution for cross-modality biomarker translation—rather than as confirmation, and we make no causal-ordering claim between glial and neuronal events, the dominant model placing reactive astrocytes downstream of activated microglia.

These results sit alongside, rather than against, the recent generation of trajectory-based AD atlases. Large single-nucleus studies increasingly reconstruct disease progression from cellular composition and community dynamics rather than from individual marker boundaries: integrated multimodal atlases order donors along a continuous pseudo-progression score[4], multiregion dissection resolves regionally staggered cellular responses[5,44], and community-level modelling of the aged prefrontal cortex has identified two distinct ageing trajectories—one progressing to AD and one reflecting alternative brain ageing—together with specific glial and neuronal subpopulations, including an astrocyte state that mediates the effect of tau pathology on cognitive decline[45]. Our contribution is complementary rather than competing: instead of proposing a further trajectory or cell state, we ask whether a marker-defined transition boundary localized in one region behaves as a brain-wide event, and we provide the permutation and cross-dataset controls required to answer that question. The attenuation of the MTG marker boundary in prefrontal cortex is thus not in tension with trajectory structure being recoverable there from composition: marker-average shifts and cell-community abundance are different readouts that need not coincide, and treating them as interchangeable is exactly the inference the framework is built to flag.

The region- and cohort-dependence we document is consistent, too, with current understanding of astrocyte heterogeneity. Subtype-resolved single-nucleus analyses show that astrocytes and oligodendrocytes undergo distinct, subtype-specific transcriptional changes in AD, with particular reactive subtypes concentrated at specific anatomical locations[46], and that disease-associated programs are only partially shared across cohorts[18,21]. A marker panel held fixed across regions will therefore sample different mixtures of these subtypes in different tissues, so a boundary defined by their aggregate behaviour need not transfer even when the underlying biology is conserved at finer resolution. This offers one mechanistic reading of why the HMOX1-led MTG signature inverts in entorhinal cortex while a within-region reactive positive control behaves as expected, and of why oligodendrocytes—whose AD-associated changes are themselves subtype-specific[46]—were the single MTG population in which the transition was not recovered. Two non-exclusive mechanisms therefore plausibly underlie boundary failure across regions: regionally staggered disease trajectories that place equivalent cellular events at different pseudo-progression positions, and region-specific differences in astrocyte subtype mixture, which we confirm directly (the MTG-enriched Astro_1 supertype is ∼3.8-fold depleted in A9 relative to MTG)—compounded at the analytic level by global transcriptional-amplitude scaling that can masquerade as coordinated module change. None of these requires a failure of the underlying biology; each instead limits the tissue and scale at which a single boundary projection remains valid.

Several limitations apply. The entorhinal-cortex analysis relies on a small, demographically homogeneous cohort (10 male ApoE3/3 donors) and is best supported by its proxy-independent reactive-astrocyte positive control, which confirms the cohort is able to detect a coordinated reactive signal and so makes the divergent result unlikely to be a power artefact, although larger multi-donor and multi-ancestry cohorts will be needed to confirm the entorhinal divergence; single-nucleus sequencing undercounts low-abundance transcripts; cross-sectional pseudo- progression cannot establish within-subject temporal ordering; the cerebrospinal-fluid comparison is limited to the 21 of 44 panel markers detectable by mass spectrometry, so intracellular drivers central to the brain signal (HMOX1, FKBP5, STAT3) could not be tested in that compartment; the generalizability framework itself has so far been exercised on a limited set of regions, cohorts and a single companion marker, so its capacity to separate conserved from dataset-specific signals is demonstrated here rather than exhaustively validated; and the boundaries carry no prognostic value beyond cognition and amyloid–tau status. Prognostic prediction was not an aim of this framework: the boundaries are cross-sectional staging constructs whose purpose is to mark where a molecular transition is reproducible, not to forecast decline. In conclusion, defining the regional, cohort and modality limits of a candidate molecular transition—and separating genuine conservation from global-expression artefact—provides a more reliable foundation for molecular staging in Alzheimer’s disease than a single-region claim generalized prematurely to the whole brain, and the framework is directly applicable to other progressive proteinopathies in which trajectory boundaries are increasingly used for staging and stratification. In particular, by certifying astrocytic PTGDS as a Class II anchor that passes every reproducibility test while a larger single-dataset signal does not, the framework demonstrates its own discriminating power, establishing a controlled and generalizable foundation for molecular staging that stands on its own independently of any single downstream application.

### Limitations of the study

Several limitations qualify these conclusions. First, the audit certifies reproducibility, not biological causation: a boundary that passes all four axes is a more credible staging anchor but is not thereby shown to drive pathology. Second, the cross-modality and cross-cohort axes are bounded by the available data—cerebrospinal-fluid proteomes detect only a subset of the marker panel, and the independent entorhinal cohort is small (n = 10), so a negative cross-cohort result constrains rather than refutes a regional claim. Third, the synthetic benchmark fixes ground truth by construction; its operating characteristics (sensitivity 100%, specificity 94%) describe the planted regimes and the discrete bin grid used here and need not transfer unchanged to every real dataset. Fourth, the framework localizes and certifies boundaries but does not, on its own, establish their prognostic value—indeed, the boundary examined here carried none beyond baseline cognition and amyloid–tau status. Finally, this is a secondary, cross-sectional re-analysis of existing atlases; longitudinal, within-individual validation of the certified boundaries remains future work.

## STAR★Methods

### Key resources table

| REAGENT or RESOURCE | SOURCE | IDENTIFIER |
| --- | --- | --- |
| *Deposited data* |  |  |
| SEA-AD snRNA-seq (MTG, DLPFC) | Allen Institute | portal.brain-map.org |
| Entorhinal cortex snRNA-seq | Leng et al.[29] | GEO: GSE147528 |
| ADNI CSF proteomics (TMT-MS; SomaScan) | ADNI | adni.loni.usc.edu |
| Analysis code and bin-level data (this paper) | This paper | github.com/YoungOukKim/AD-Boundary-Audit<br>Zenodo:<br>https://doi.org/10.5281/zenodo.21079773 |
| *Software and algorithms* |  |  |
| R (v4.3) | R Core Team | r-project.org |
| segmented; cluster; edgeR | CRAN; Bioconductor | cran.r-project.org; bioconductor.org |
| Boundary-detection engine and synthetic benchmark | This paper | github.com/YoungOukKim/AD-Boundary-Audit<br>Zenodo:<br>https://doi.org/10.5281/zenodo.21079773 |

### Resource availability

#### Lead contact

Further information and requests for resources should be directed to and will be fulfilled by the lead contact, YoungOuk Kim.

#### Materials availability

This study is a computational re-analysis of existing data and did not generate new unique reagents or materials.

#### Data and code availability

All analysis code and the bin-level data that deterministically regenerate every figure, table and reported statistic (global seed 42; single-command run_all) are publicly available at https://github.com/YoungOukKim/AD-Boundary-Audit, including R/analysis/synthetic_benchmark.R, which reproduces the synthetic ground-truth benchmark (Figure 7) end-to-end under the same seed; the repository DOI is listed in the key resources table. This paper analyses previously published data: SEA-AD snRNA-seq are available from the Allen Institute portal; the entorhinal cohort from GEO (GSE147528); ADNI proteomic, biomarker and cognitive data upon application to the ADNI Data and Publications Committee. Any additional information required to reanalyse the data reported in this paper is available from the lead contact upon request.

### Method details

#### Datasets

SEA-AD middle temporal gyrus single-nucleus RNA-sequencing data (1.3 million nuclei, 84 donors) were downloaded from the Allen Institute portal (SEAAD_MTG_RNAseq_final- nuclei.2024-02-13.h5ad). Subclass labels, the continuous pseudo-progression score (CPS), and donor metadata were extracted from the h5ad /obs/ group. Astrocyte nuclei were identified by Subclass labels matching ^Astro and filtered to CPS ≥ 0.1, yielding 67,419 astrocytes. SEA-AD dorsolateral prefrontal cortex (Brodmann area 9) snRNA-seq was processed identically; where CPS was absent from A9 /obs/, it was joined from MTG by Donor ID. The independent entorhinal cortex (EC) cohort (Leng et al. 2021; GSE147528; 10 male ApoE3/3 donors) was obtained as raw 10x H5 matrices. ADNI CSF proteomic data (SomaScan 7K, post-QC 20230620 release) were accessed via the ADNI portal; clinical metadata (DXSUM, MMSE, MOCA, PTDEMOG, APOERES) were loaded from ADNIMERGE2[47]. The Emory TMT-MS CSF cohort (n = 1,104 of the 1,105-donor Emory cohort) was used for robustness and prognostic analyses.

#### Marker panel design

The 44-marker reactive-astrocyte panel spanned A1/A2 reactive signatures, inflammation, vascular endothelium, AD risk genes, iron/oxidative stress, neurotrophic factors and core AD proteins (Table S1), and deliberately excluded PTGDS, LCN2 and MAPT to enable independent boundary detection. MCT4 (SLC16A3) was excluded from the panel because it is the focus of a companion astrocyte-neuron lactate-shuttle study. For panel-sensitivity analysis, a 39-gene panel including PTGDS and LCN2 was constructed. A separate 21-gene metabolic-supply set (SLC16A1/3/7, LDHA/B, SLC2A1/3, glycolytic enzymes, NDUFS1, myelin genes MOG/MAG/MBP/PLP1, MAPT, TFEB) was analysed to test metabolic conservation.

#### Data preprocessing and boundary detection

For SEA-AD, panel-gene expression was read from the primary expression matrix (.X) of the h5ad file via rhdf5[48]; matrices detected as raw integer counts were transformed as log(1 + x) and matrices already on a log scale were used as provided, after which values were z-scored per gene within each cell type. Astrocytes were partitioned into nine CPS bins of width 0.1 (bins < 50 cells excluded), and within each bin the mean z-score per marker was computed. For EC, raw 10x matrices were processed with emptyDrops cell calling, per-sample layer joining, log- normalisation, Harmony integration and clustering; donor pseudobulk profiles per cell type were tested with edgeR; donor-level (pseudobulk) aggregation was used for all cohort contrasts to avoid the false-discovery inflation and highly-expressed-gene bias of cell-level differential-expression testing. A within-cohort positive control contrasted GFAP-high versus GFAP-low astrocytes. A separate pseudoreplication control compared cell-level (Wilcoxon, nuclei as replicates) with donor-level (edgeR pseudobulk) differential expression across all SEA-AD MTG cell types over a fixed 4,000-gene random universe, quantifying DEG-count inflation, its dependence on nucleus number, and expression-level bias (Figure S4). Nine algorithms were applied to each cohort’s bin-aggregated matrix: piecewise linear regression on PC1–PC3, CUSUM, Mahalanobis maximum-gap, Ward.D2 hierarchical clustering (k = 3), k-means (k = 3, seed 42), dynamic-programming optimal change-point, variance-jump, trajectory inflection, and segmented regression with the Davies test[49,50]. Boundary calls were aggregated across the 19 algorithm–component combinations.

##### Reproducibility

The consensus engine is deterministic (global seed 42); re-running the nine algorithms on the committed bin-level matrices reproduces every clustering, partitioning, change-point, Mahalanobis, variance-jump and trajectory-inflection call exactly. Segmented-regression breakpoints on principal components lacking a significant Davies signal depend on the segmented implementation and its bootstrap-restart setting, so we fix n.boot = 0 and record package versions to make the committed breakpoints reproduce deterministically; all boundaries reported here—the CPS-0.21 transition and the CPS-0.39–0.42 secondary (dynamic-programming change-point and the ADNI MMSE-26 anchor)—are recovered independently of this setting. Analysis code and bin-level data that regenerate every figure and table are provided in the repository (Data and code availability).

#### Permutation testing and global-expression correction

For each cohort and panel, 1,000 random panels of matched size were drawn from the full feature space, the bin-aggregated matrix recomputed, and empirical p-values for each discovered boundary obtained (±0.05 CPS or ±1.0 MMSE tolerance). For the metabolic analysis, per-cell-type global-expression correction was applied before conservation classification by subtracting, within each region and cell type, the mean effect across all genes from each gene’s effect; a gene was classified conserved or divergent only if its corrected effect exceeded |0.10| in both regions. As a negative control for this correction, 14 constitutive housekeeping genes (ACTB, GAPDH, B2M, PGK1, TBP, RPL13A, HPRT1, PPIA, YWHAZ, SDHA, UBC, RPLP0, GUSB, POLR2A) were carried through the identical per-cell-type, per-CPS-bin standardization, and the mean absolute binned z-score is reported before and after correction. Astrocyte subtype composition was compared between MTG and A9 from per-donor proportions of the six SEA-AD astrocyte supertypes, centred-log-ratio transformed and tested by Wilcoxon rank-sum with Benjamini– Hochberg correction (n = 84 donors). Because the A9 single-nucleus object does not carry a per-nucleus pseudo-progression score, donor-level CPS values were mapped from MTG by donor identity for any CPS-stratified A9 analysis; CPS is a donor-level pathological coordinate rather than a region-intrinsic trajectory, so this assigns each donor a single disease-stage value and does not transplant an MTG-derived progression axis into A9.

#### Synthetic benchmark with planted ground-truth boundaries

To establish the audit’s operating characteristics on data with known ground truth, we constructed synthetic marker × CPS-bin matrices in which the presence and location of a transition boundary are set by design. Each dataset comprised a 3,000-gene background—55% flat noise and 45% logistic transitions with inflection points drawn uniformly over the late trajectory (CPS 0.48–0.84)—calibrated so that random panels reproduce the empirical null shape seen in real data (change-points concentrated late and sparse early; in the real MTG astrocyte data 0/1,000 random panels recovered the early boundary while ≈40% fell near CPS 0.54). From each background we drew 44-marker panels (matching the real panel size) under five ground-truth regimes: (i) a real panel-specific boundary, a co-regulated step at CPS 0.20 below the null change-point floor; (ii) a global-expression offset, the same step shared by every background gene; (iii) a null panel with no coordinated transition; (iv) a monotonic ramp; and (v) a single-marker decoy in which one marker carries a large step against an otherwise structureless panel. The identical detection engine (nine algorithms, 19 components, seed 42) and audit gates were then applied. The three gates were fixed a priori from the null rather than tuned to the answer: a boundary was certified only if its algorithmic-consensus support reached the 95th percentile of the random-panel null support (S* = 6; nonparametric bootstrap 95% CI 5–7), its random-panel specificity P ≤ 0.05 (NPERM = 300 panels), and no single marker shifted the boundary by more than one bin (≤ 0.05 CPS) on leave-one-marker-out removal. The benchmark used a finer fixed grid (20 bins, 300 random panels) than the cell-count-limited real analysis (nine CPS bins, 1,000 panels); the detection engine is bin-count-agnostic and the null shape is stable across both.

Across 24 replicate panels per regime, the audit certified every real panel-specific boundary (sensitivity 100%) and rejected all four artefact classes (specificity 94%; global offset 100%, null 92%, ramp 92%, decoy 92%), each excluded primarily by a distinct axis: the global offset passed consensus support but was reproduced by random panels and failed specificity (median P = 0.21); null and ramp panels failed support; and the single-marker decoy passed support and specificity yet was flagged by leave-one-marker-out in 22 of 24 replicates (median maximum single-marker boundary shift 0.30 CPS versus 0.00 for real panels; the two that slipped had boundaries that, by chance, shifted only one bin on driver removal). No single algorithm localized the planted boundary reliably—per-method localization error ranged from 0.00 CPS (Mahalanobis, dynamic programming) to 0.60 CPS (variance-jump)—whereas the consensus error was 0.025 CPS and remained 0.022–0.025 CPS when any one of the nine algorithm families was removed, indicating that localization does not depend on the exact algorithm set. The benchmark code reproducing these results is provided (Data and code availability; Figure 7).

#### AD-GWAS convergence test

A set of 41 AD-GWAS risk genes detected in the data was specified before analysis from genome-wide significant Alzheimer’s disease loci[27,28]. For each cell type, the boundary effect of every gene was the difference in its mean z-score between nuclei above and below CPS 0.207, and the test statistic was the mean absolute boundary effect across the risk-gene set. The empirical null comprised 1,000 random gene sets drawn from a 4,000-gene universe, matched to the risk-gene set by expression decile to control for the highly-expressed-gene bias; the empirical p-value was the fraction of null sets with a statistic at least as large. Because significance derives from permutation across matched gene sets rather than from treating nuclei as independent replicates, this test is not subject to the pseudoreplication inflation controlled for elsewhere.

#### CSF marker-level association analysis

For direct marker-level external validation, baseline CSF abundances for panel markers were extracted from the ADNI SomaScan 7K matrix (earliest measurement per RID; n = 735; the HMOX1−cognition association uses this set minus HMOX1 missing values, n = 724, NA rate 1.5%, and the CSF trajectory analysis uses the consensus-clustering subset, n = 343) and from the ADNI Emory TMT-MS matrix, and merged with baseline MMSE (screening/baseline visit). Log2-transformed, per-protein z-scored abundances were related to MMSE by Spearman correlation and by binning along the SEA-AD-matched MMSE intervals. HMOX1 (SomaScan SeqId 17398-55) was quantified in SomaScan (NA rate 1.5%) but was absent from the TMT-MS quantified protein set.

#### Quantification and statistical analysis

Prognostic models were fit on the Emory TMT-MS cohort (n = 1,105 with complete baseline MMSE, Roche Elecsys CSF amyloid–tau, and panel proteomics). The endpoint was time to a ≥ 3-point MMSE decline from baseline or MMSE ≤ 23, with months from baseline computed from visit dates and censoring at the last available visit (527 events). Amyloid (A+) and tau (T+) positivity used standard ADNI Elecsys cut-offs (CSF Aβ42 < 980 pg/mL; pTau181 > 24 pg/mL). A molecular boundary score was defined as the first principal component of the available CSF panel proteins (21 of the 44 markers detected in TMT-MS; intracellular drivers such as HMOX1, FKBP5 and STAT3 were below detection), standardised and oriented so that higher values correspond to lower baseline cognition. Cox proportional-hazards models (statsmodels PHReg) compared boundary positivity defined by an MMSE threshold (≤ 26) versus the continuous molecular score, the latter adjusted for baseline MMSE, A+ and T+. Robustness analyses on the Emory cohort included binning-scheme sweeps, leave-one-marker-out (28 markers), and APOE ε4 and sex stratification. Other analyses were performed in R[51] (rhdf5[48], tidyverse[52], segmented[49], edgeR, Harmony, pROC[53], survival[54], patchwork[55]). Significance thresholds were P < 0.05 for Davies tests and permutation empirical p-values, FDR < 0.05 for edgeR contrasts, and Bonferroni-adjusted thresholds for multi-boundary comparisons.

## Declarations

### Ethics

This study is a computational secondary analysis of fully de-identified, publicly available human datasets (SEA-AD; ADNI; GSE147528). Original ethics approvals and informed consent were obtained by the respective consortia.

### Author contributions

Y.O.K. conceived the study, performed all computational analyses, and wrote the manuscript. W.M.H. supervised the project. S.J.P. contributed to study design and interpretation. Y.C.K. contributed to data curation and QC. Y.E.C. contributed to data collection and interpretation. Y.O.K., W.M.H. and S.J.P. contributed equally.

## Acknowledgements

We thank the Allen Institute for Brain Science for the SEA-AD resource, and the study participants and their families. Data used in preparation of this article were obtained in part from the Alzheimer’s Disease Neuroimaging Initiative (ADNI) database (adni.loni.usc.edu). As such, the investigators within the ADNI contributed to the design and implementation of ADNI and/or provided data but did not participate in analysis or writing of this report. A complete listing of ADNI investigators is available at adni.loni.usc.edu. Data collection and sharing for this project was funded by the Alzheimer’s Disease Neuroimaging Initiative (ADNI) (National Institutes of Health Grant U01 AG024904) and DOD ADNI (Department of Defense award number W81XWH-12-2-0012). ADNI is funded by the National Institute on Aging, the National Institute of Biomedical Imaging and Bioengineering, and through generous contributions from the following: AbbVie, Alzheimer’s Association; Alzheimer’s Drug Discovery Foundation; Araclon Biotech; BioClinica, Inc.; Biogen; Bristol-Myers Squibb Company; CereSpir, Inc.; Cogstate; Eisai Inc.; Elan Pharmaceuticals, Inc.; Eli Lilly and Company; EuroImmun; F. Hoffmann -La Roche Ltd and its affiliated company Genentech, Inc.; Fujirebio; GE Healthcare; IXICO Ltd.; Janssen Alzheimer Immunotherapy Research & Development, LLC.; Johnson & Johnson Pharmaceutical Research & Development LLC.; Lumosity; Lundbeck; Merck & Co., Inc.; Meso Scale Diagnostics, LLC.; NeuroRx Research; Neurotrack Technologies; Novartis Pharmaceuticals Corporation; Pfizer Inc.; Piramal Imaging; Servier; Takeda Pharmaceutical Company; and Transition Therapeutics. The Canadian Institutes of Health Research is providing funds to support ADNI clinical sites in Canada. Private sector contributions are facilitated by the Foundation for the National Institutes of Health (www.fnih.org). The grantee organization is the Northern California Institute for Research and Education, and the study is coordinated by the Alzheimer’s Therapeutic Research Institute at the University of Southern California. ADNI data are disseminated by the Laboratory for Neuro Imaging at the University of Southern California.

## Supplemental figures

**Figure S1.**
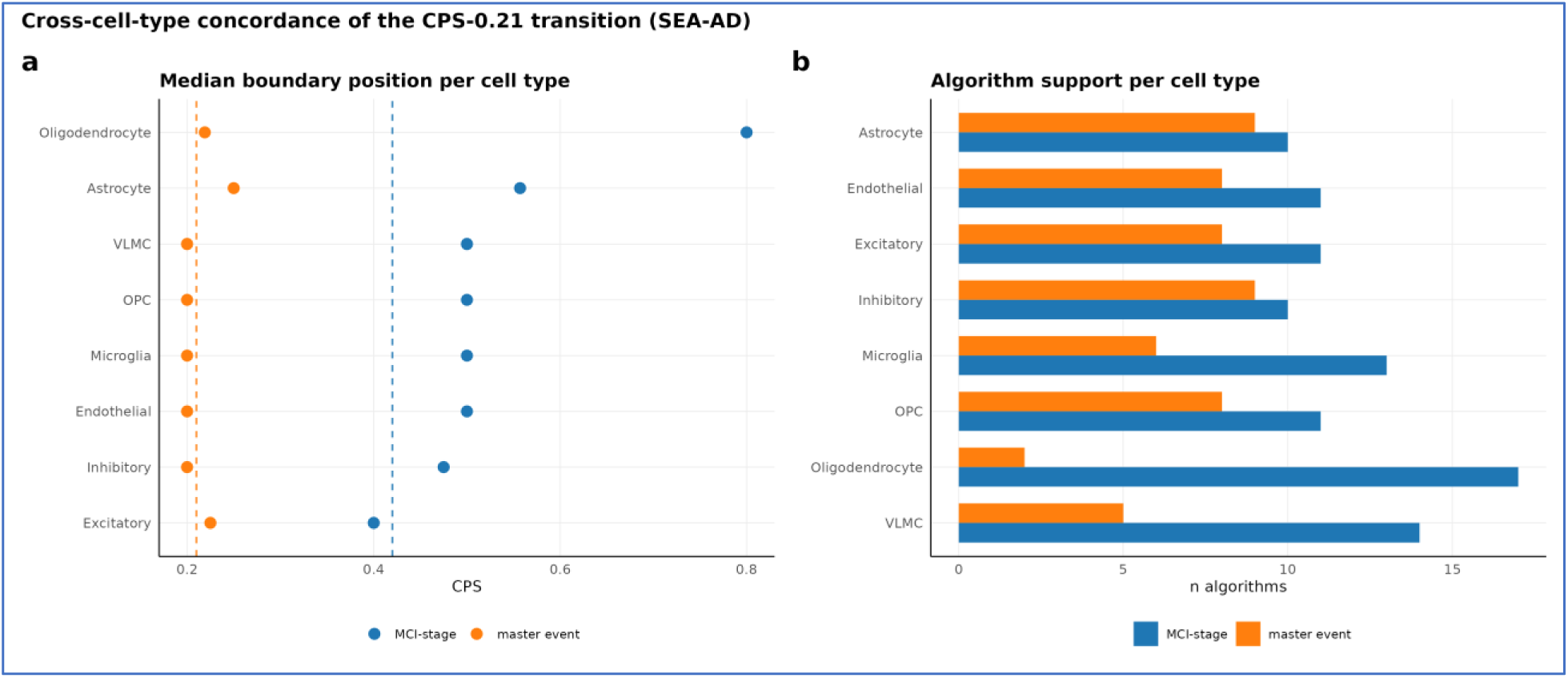
The CPS-0.21 transition is recovered across MTG cell populations. The identical 44-marker panel and consensus framework were applied to eight major MTG cell types. **(a)** Median boundary position (CPS) per cell type for this transition (CPS≈0.21) and the secondary MCI-stage transition; this transition localizes near CPS 0.21 in seven of eight populations, with oligodendrocytes the sole outlier. (b) Number of supporting algorithms (of nine) per cell type for each boundary. FKBP5 was the most broadly shared driver across cell types, whereas the HMOX1 decrement and GFAP induction remained astrocyte - specific, indicating a coordinated, cross-cell-type event within MTG rather than an astrocyte-only phenomenon.

**Figure S2.**
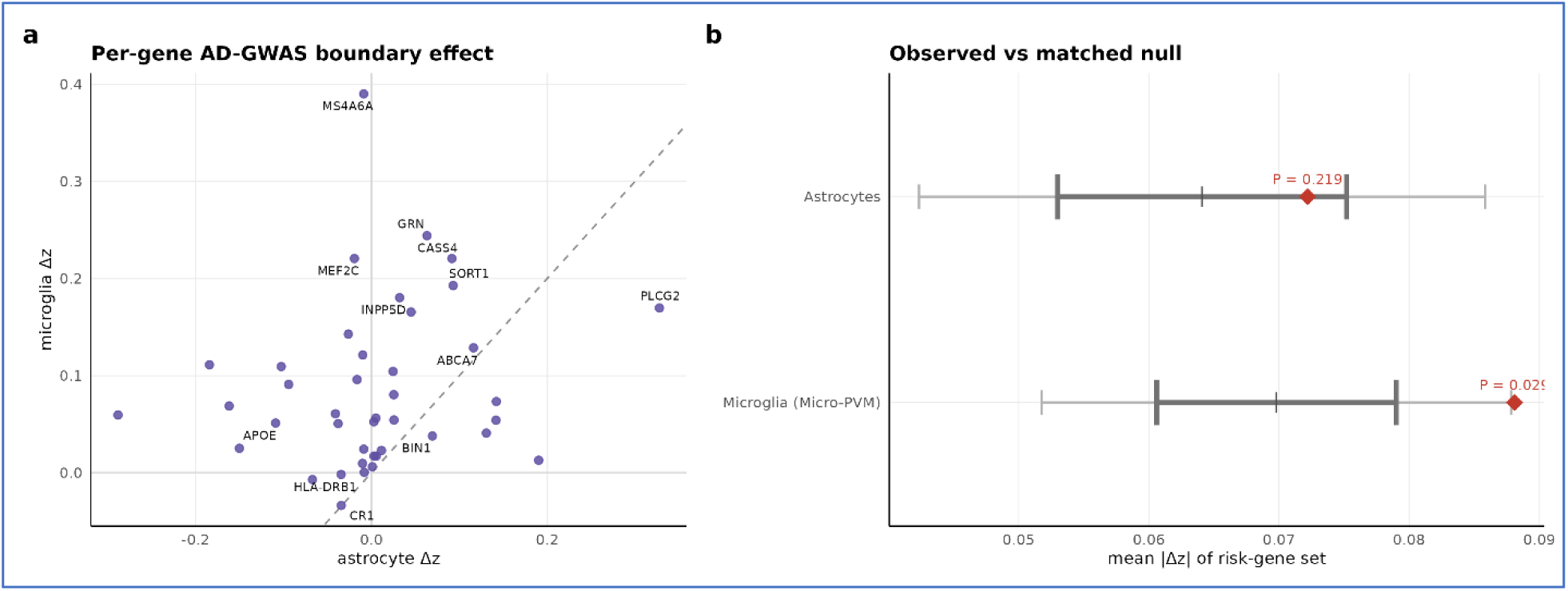
AD-GWAS risk genes converge on the CPS-0.207 boundary in microglia but not astrocytes. A pre-registered set of 41 detected AD-GWAS risk genes was tested for coordinated change across the boundary against an expression-decile-matched permutation null (1,000 random gene sets). (a) Per-gene boundary effect (Δz) in astrocytes versus microglia; most genes lie above the dashed identity line (larger in microglia) while astrocytic effects scatter around zero. (b) Observed mean |Δz| of the risk-gene set (red diamond) versus the matched null (band: mean ±1 and ±1.96 s.d.): astrocytes P = 0.22, microglia P = 0.029 (nominal; ≈ 0.06 after correction for the two cell types tested).

**Figure S3.**
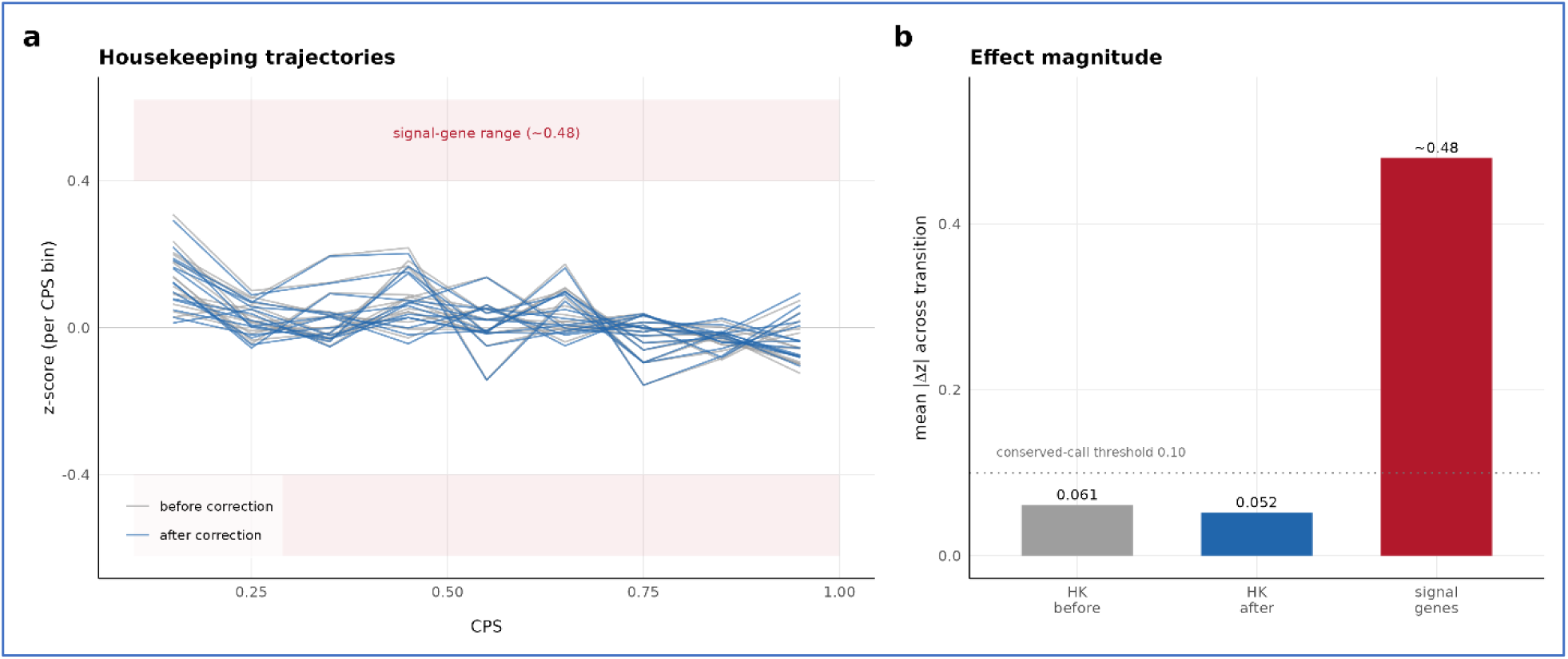
Housekeeping negative control for the global-expression correction. **(a)** Trajectories of the 14 constitutive housekeeping genes across CPS before (grey) and after (blue) the genome-wide correction remain flat near zero, far from the signal-gene range (|Δz| ≈ 0.48). **(b)** Mean |Δz| across the transition for the housekeeping panel before and after correction (0.061 and 0.052) is well below both the conserved- call threshold (0.10) and the signal-gene scale (∼0.48), confirming that the correction removes only the shared global offset and does not manufacture gene-specific signal. Astrocytes, MTG; generated by R/figures/SuppFig_global_correction_control.R.

**Figure S4.**
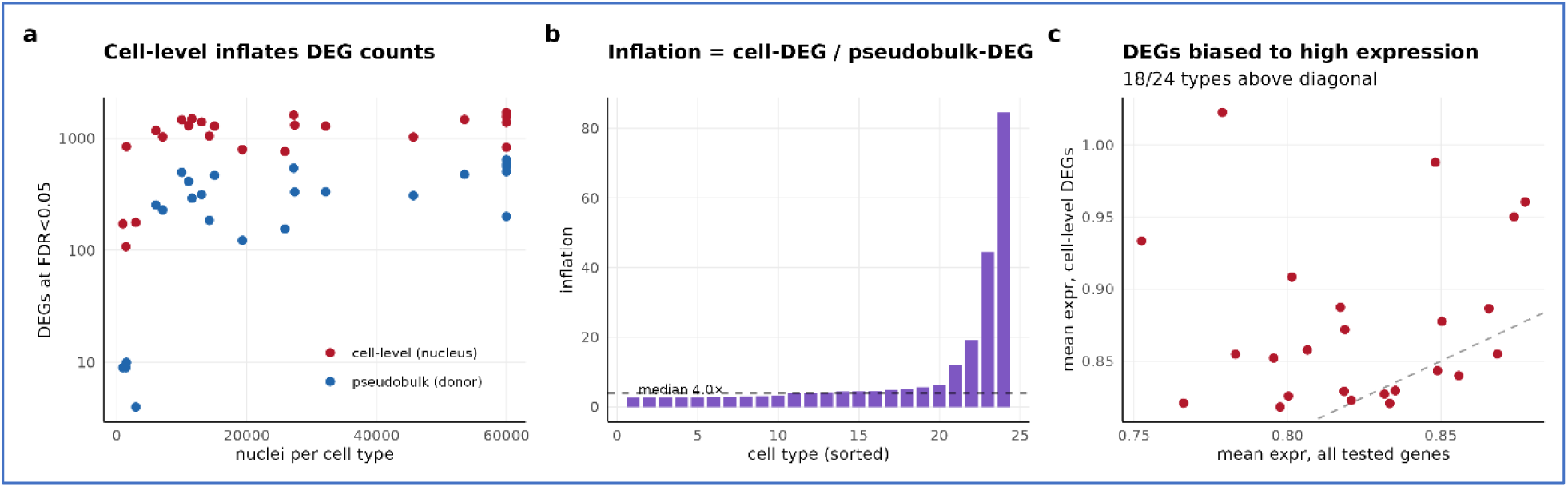
Donor-level (pseudobulk) testing is required. Across 24 SEA-AD MTG cell types (4,000- gene random universe): **(a)** genes called differentially expressed at FDR<0.05 by cell-level (nucleus-as- replicate) versus donor-level pseudobulk (edgeR); cell-level counts scale with nucleus number (r = 0.52). **(b)** Per-cell-type inflation (cell-level / pseudobulk DEGs): median 4.0×, up to ∼85× in cell types with few donors. **(c)** Cell-level DEGs are enriched for highly expressed genes (18 of 24 cell types above the diagonal).

**Figure S5.**
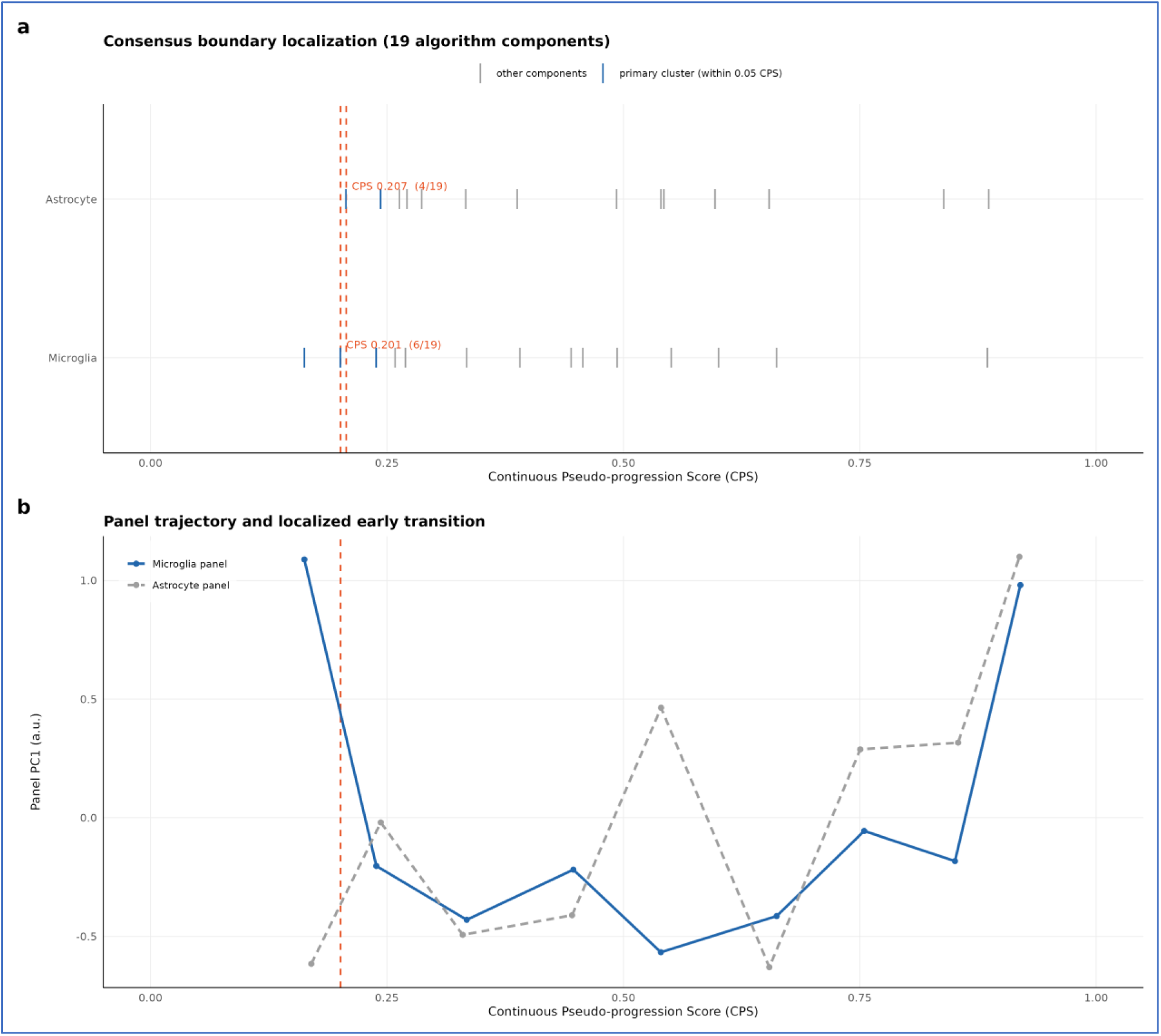
The audit generalizes across cell types. The identical multi-algorithm consensus audit, applied to an independent microglia-specific 44-marker panel, localizes a transition concordant with the reactive-astrocyte panel. **(a)** Boundary localization across the 19 algorithm components for the astrocyte and microglia panels; the primary cluster (within 0.05 CPS of the Ward boundary) localizes at CPS 0.207 (astrocyte) and CPS 0.201 (microglia). **(b)** Panel PC1 trajectory across CPS for the microglia panel (astrocyte panel shown for reference), with the localized early transition marked. Generated from bundled bin-level data by R/figures/SuppFig_celltype_generalization.R.

## Supplemental tables

**Table S1.** Composition of the 44-marker reactive-astrocyte panel.

| Category | Markers |
| --- | --- |
| Reactive astrocyte | GFAP, SERPINA3, VIM, FBLN5, AQP4, MERTK, MFGE8, SOX9, FKBP5 |
| A1 signaling | C3, STAT1, STAT3, SOCS3, NFKB1 |
| Inflammation | IL1B, IL6, TNF, CCL2, CXCL12, CXCL16 |
| Vascular | ICAM1, VCAM1, VWF, SELE, OCLN, PECAM1 |
| AD risk | APOE, CLU, TREM2, CR1, CD33 |
| Iron / oxidative | FTH1, FTL, HMOX1, NFE2L2, TFRC |
| Injury / protective | NEFL, BDNF, GDNF, NGF |
| AD core / cell death | APP, CASP3, BCL2, TP53 |

*The panel was fixed a priori and held constant across all cohorts and cell types, spanning 8 functional categories (44 markers). PTGDS, LCN2 and MAPT were deliberately excluded from the panel to permit independent boundary detection and to keep diagnostic markers separate from boundary-detection markers; MCT4 (SLC16A3) was excluded as the focus of a companion lactate-shuttle study.*

**Table S2.**
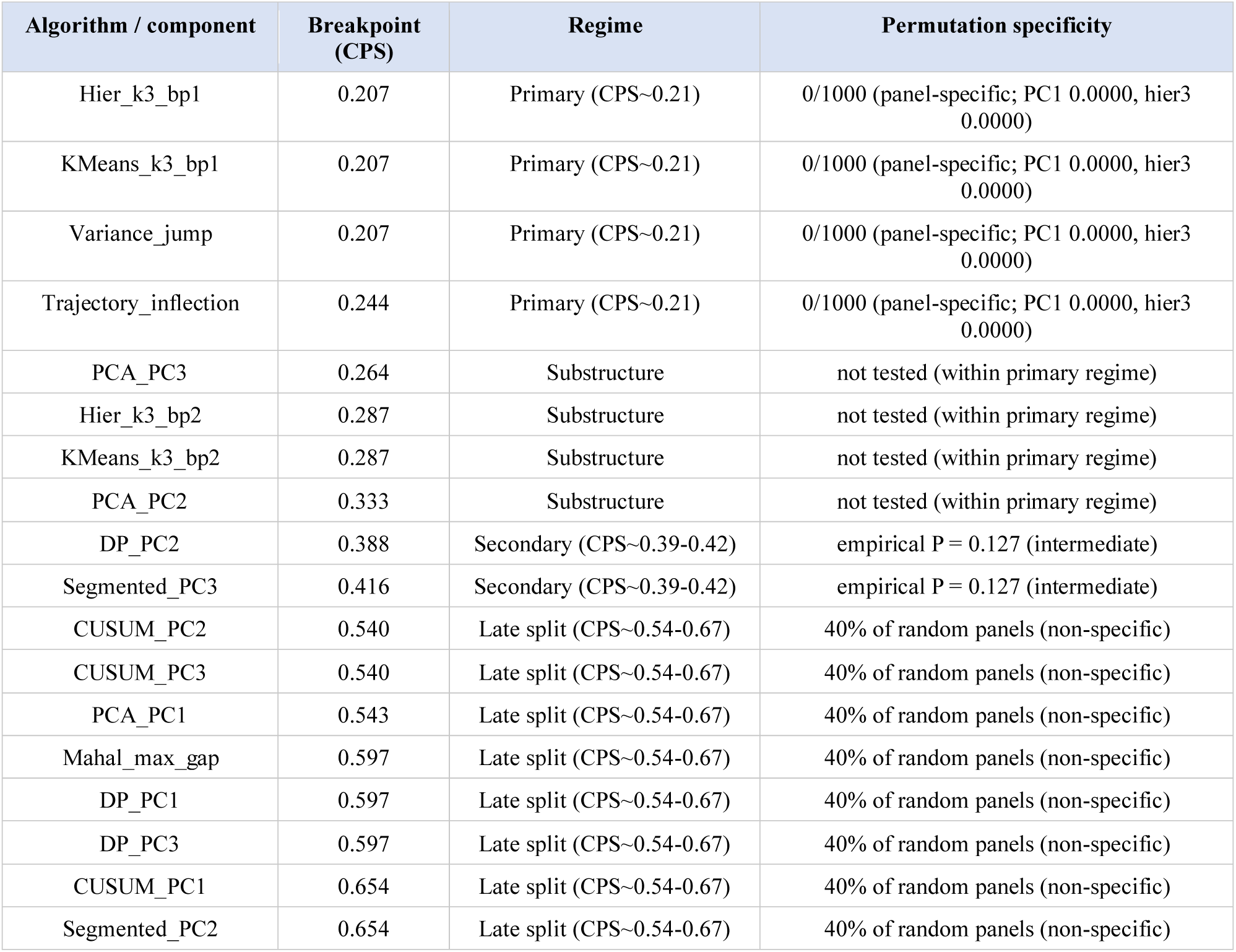
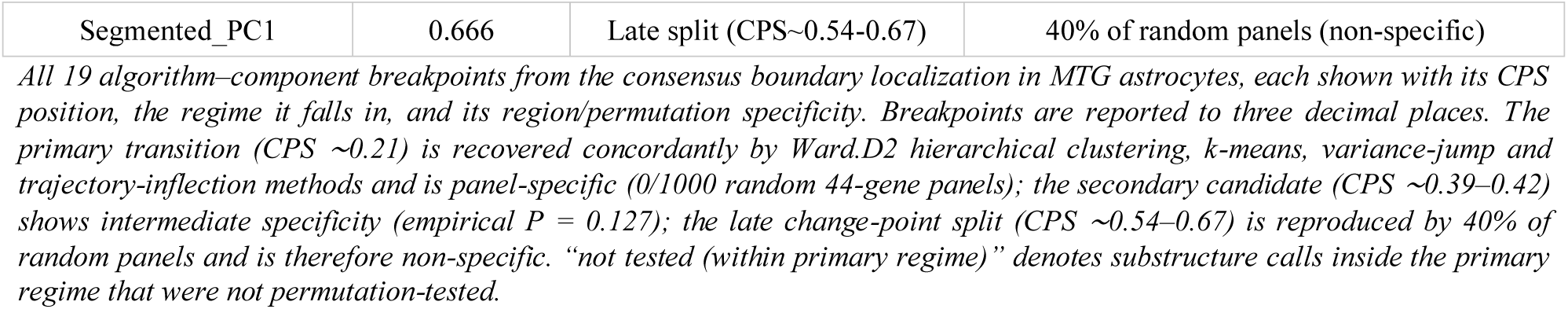
Full per-algorithm boundary distribution across the consensus localization.

| Algorithm / component | Breakpoint (CPS) | Regime | Permutation specificity |
| --- | --- | --- | --- |
| Hier_k3_bp1 | 0.207 | Primary (CPS~0.21) | 0/1000 (panel-specific; PC1 0.0000, hier3 0.0000) |
| KMeans_k3_bp1 | 0.207 | Primary (CPS~0.21) | 0/1000 (panel-specific; PC1 0.0000, hier3 0.0000) |
| Variance_jump | 0.207 | Primary (CPS~0.21) | 0/1000 (panel-specific; PC1 0.0000, hier3 0.0000) |
| Trajectory_inflection | 0.244 | Primary (CPS~0.21) | 0/1000 (panel-specific; PC1 0.0000, hier3 0.0000) |
| PCA_PC3 | 0.264 | Substructure | not tested (within primary regime) |
| Hier_k3_bp2 | 0.287 | Substructure | not tested (within primary regime) |
| KMeans_k3_bp2 | 0.287 | Substructure | not tested (within primary regime) |
| PCA_PC2 | 0.333 | Substructure | not tested (within primary regime) |
| DP_PC2 | 0.388 | Secondary (CPS~0.39-0.42) | empirical P = 0.127 (intermediate) |
| Segmented_PC3 | 0.416 | Secondary (CPS~0.39-0.42) | empirical P = 0.127 (intermediate) |
| CUSUM_PC2 | 0.540 | Late split (CPS~0.54-0.67) | 40% of random panels (non-specific) |
| CUSUM_PC3 | 0.540 | Late split (CPS~0.54-0.67) | 40% of random panels (non-specific) |
| PCA_PC1 | 0.543 | Late split (CPS~0.54-0.67) | 40% of random panels (non-specific) |
| Mahal_max_gap | 0.597 | Late split (CPS~0.54-0.67) | 40% of random panels (non-specific) |
| DP_PC1 | 0.597 | Late split (CPS~0.54-0.67) | 40% of random panels (non-specific) |
| DP_PC3 | 0.597 | Late split (CPS~0.54-0.67) | 40% of random panels (non-specific) |
| CUSUM_PC1 | 0.654 | Late split (CPS~0.54-0.67) | 40% of random panels (non-specific) |
| Segmented_PC2 | 0.654 | Late split (CPS~0.54-0.67) | 40% of random panels (non-specific) |
| Segmented_PC1 | 0.666 | Late split (CPS~0.54-0.67) | 40% of random panels (non-specific) |

*All 19 algorithm–component breakpoints from the consensus boundary localization in MTG astrocytes, each shown with its CPS position, the regime it falls in, and its region/permutation specificity. Breakpoints are reported to three decimal places. The primary transition (CPS ~0.21) is recovered concordantly by Ward.D2 hierarchical clustering, k-means, variance-jump and trajectory-inflection methods and is panel-specific (0/1000 random 44-gene panels); the secondary candidate (CPS ~0.39–0.42) shows intermediate specificity (empirical $P = 0.127$ ); the late change-point split (CPS ~0.54–0.67) is reproduced by 40% of random panels and is therefore non-specific. “not tested (within primary regime)” denotes substructure calls inside the primary regime that were not permutation-tested.*

**Table S3.**
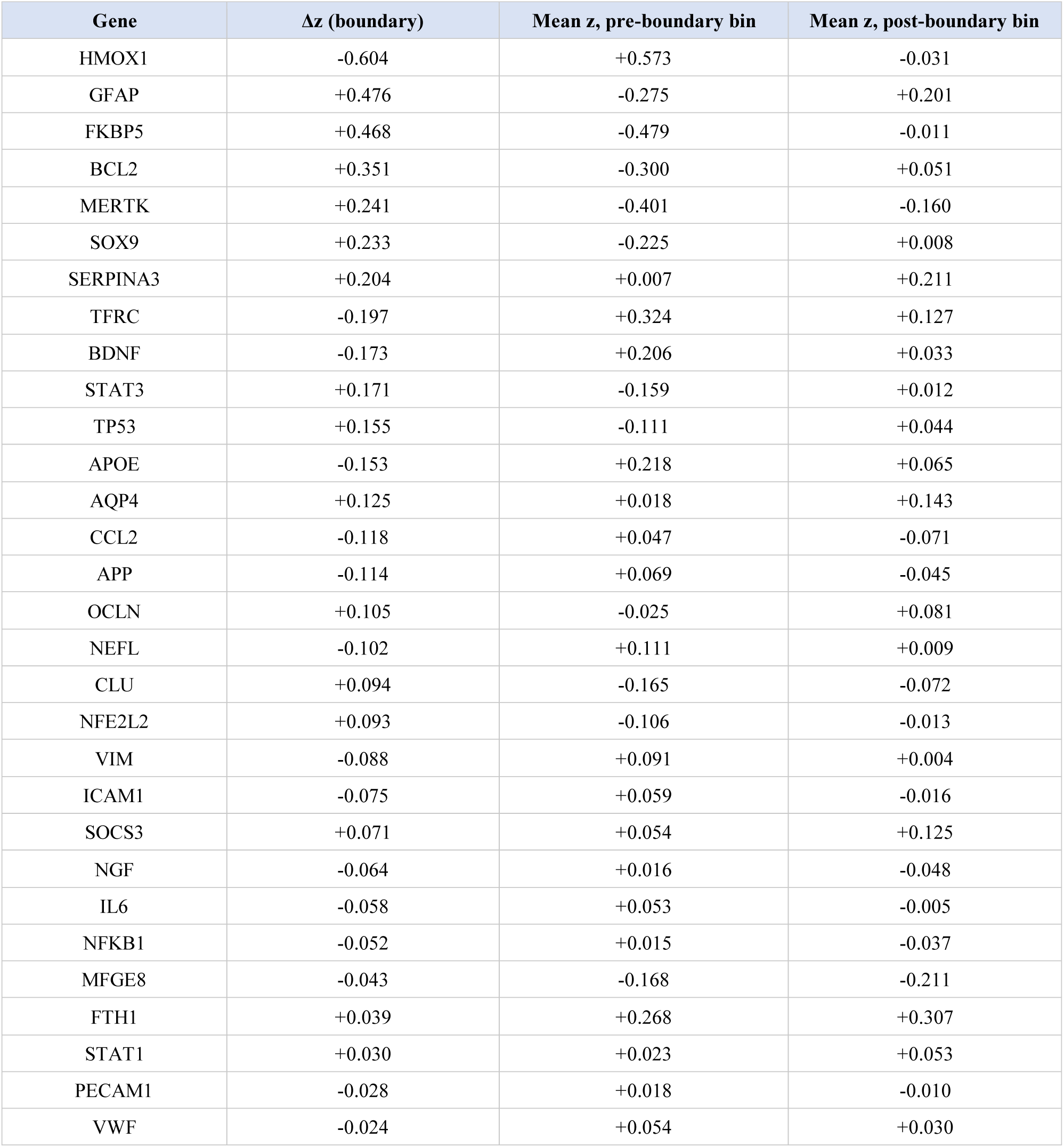
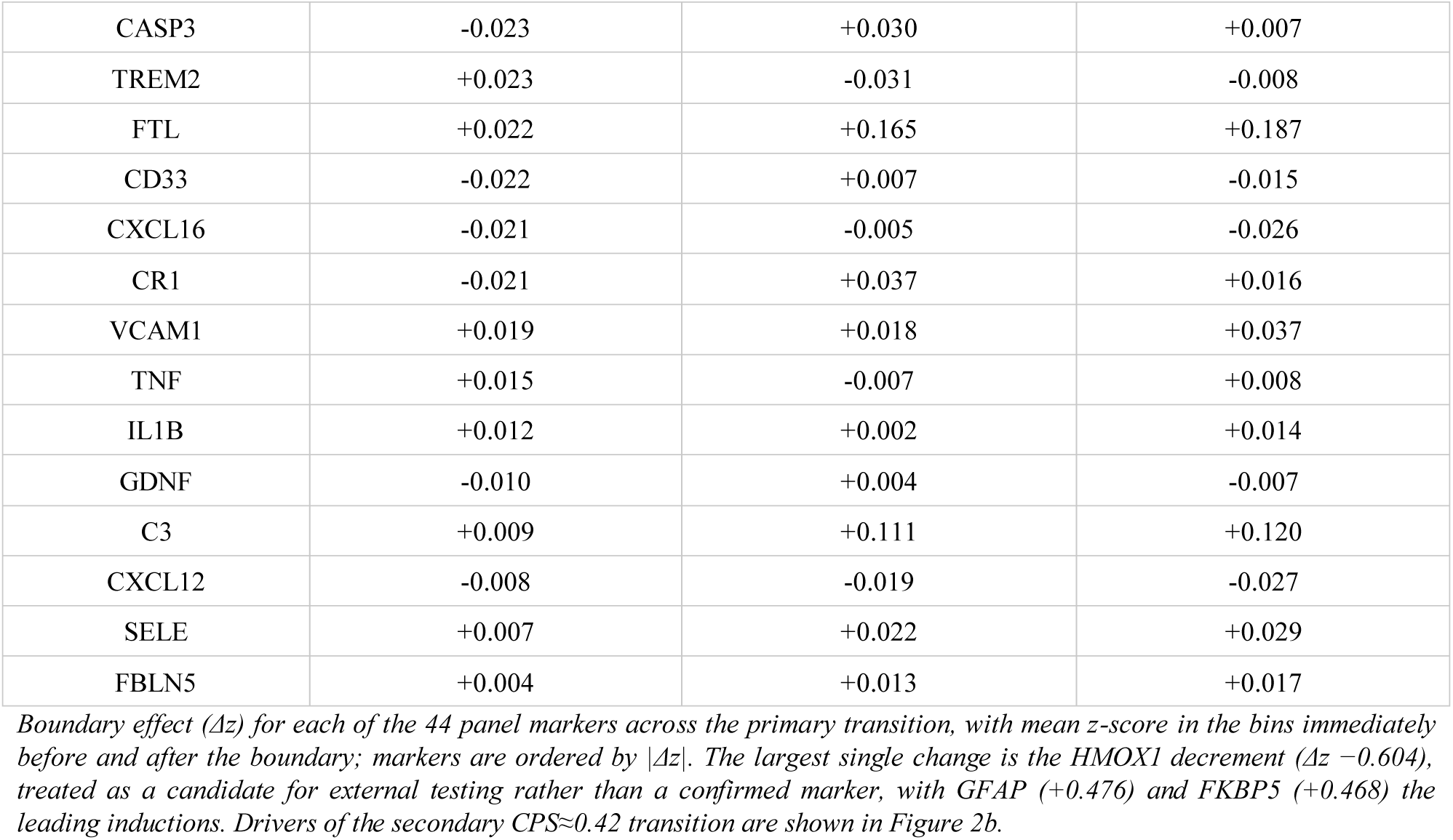
Per-marker drivers of the primary CPS≈0.21 transition in MTG astrocytes.

**Table S4.** Boundary reproducibility audit across datasets.

| Dataset (modality) | Boundary detected | Leading marker (HMOX1) | Reproduced? |
| --- | --- | --- | --- |
| SEA-AD MTG (snRNA) | Yes — CPS ≈ 0.21 (perm P < 0.001) | Strong ($\Delta z - 0.60$) | Discovery |
| SEA-AD A9, prefrontal (snRNA) | Attenuated — 1/8 cell types | $\Delta z \approx - 0.06$ | No |
| Leng EC, independent (snRNA, n = 10) | Not recovered — panel reversed | Floor; FTH1 + 0.54 (opposite) | No |
| ADNI CSF (TMT-MS / SomaScan) | Partial — MMSE-aligned, discordant | Undetected / reversed ($\rho - 0.11$) | No |

Summary of whether the CPS-0.21 transition and its leading marker (HMOX1) reproduce in each dataset. snRNA, single-nucleus RNA-seq; CSF, cerebrospinal fluid; perm, permutation; $\Delta z$ , change in mean z-score across the boundary; $\rho$ , Spearman correlation with cognition.

**Table S5.**
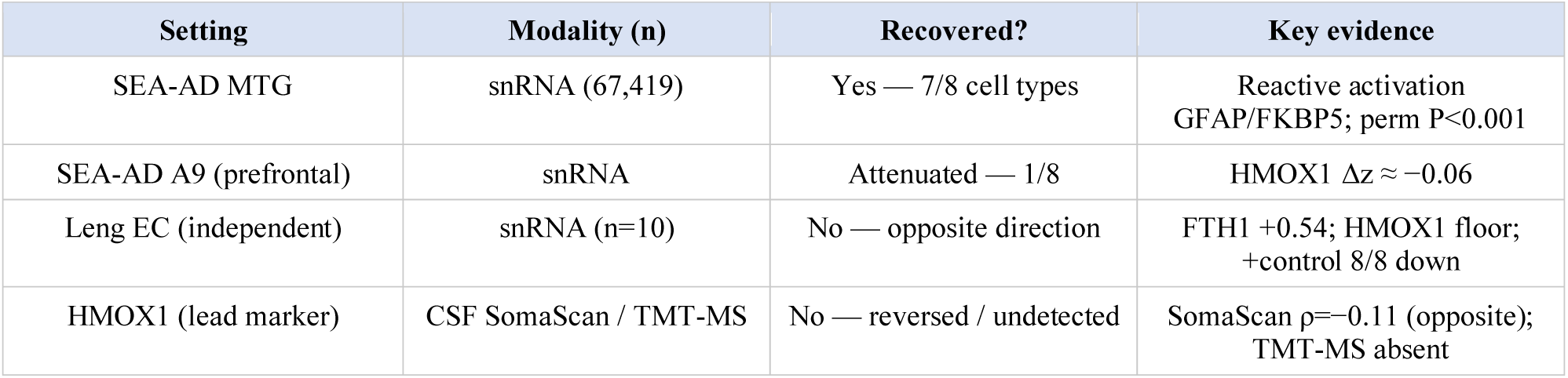
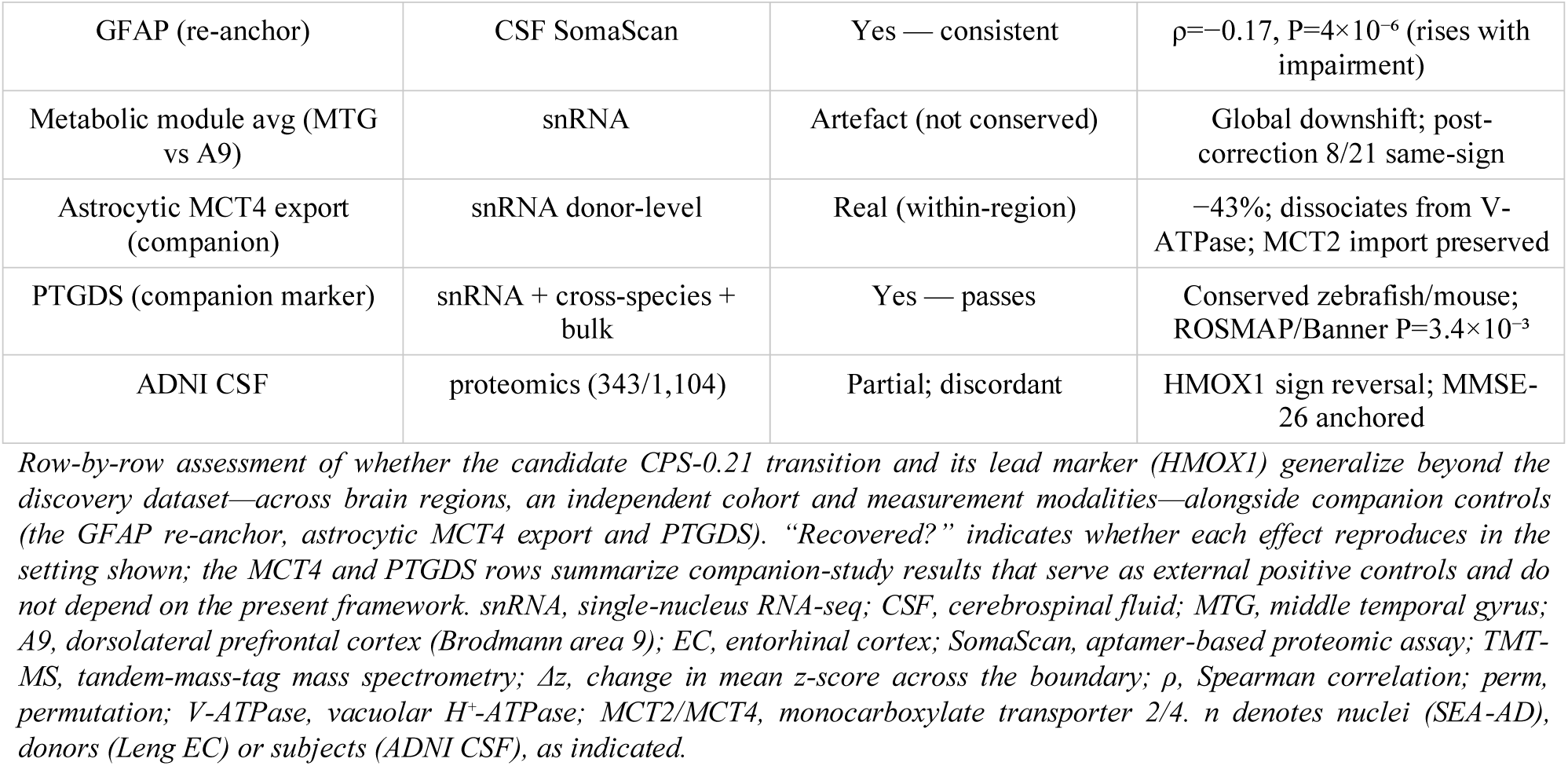
Generalization of the candidate transition and its lead marker across regions, cohorts and modalities.

**Table S6.**
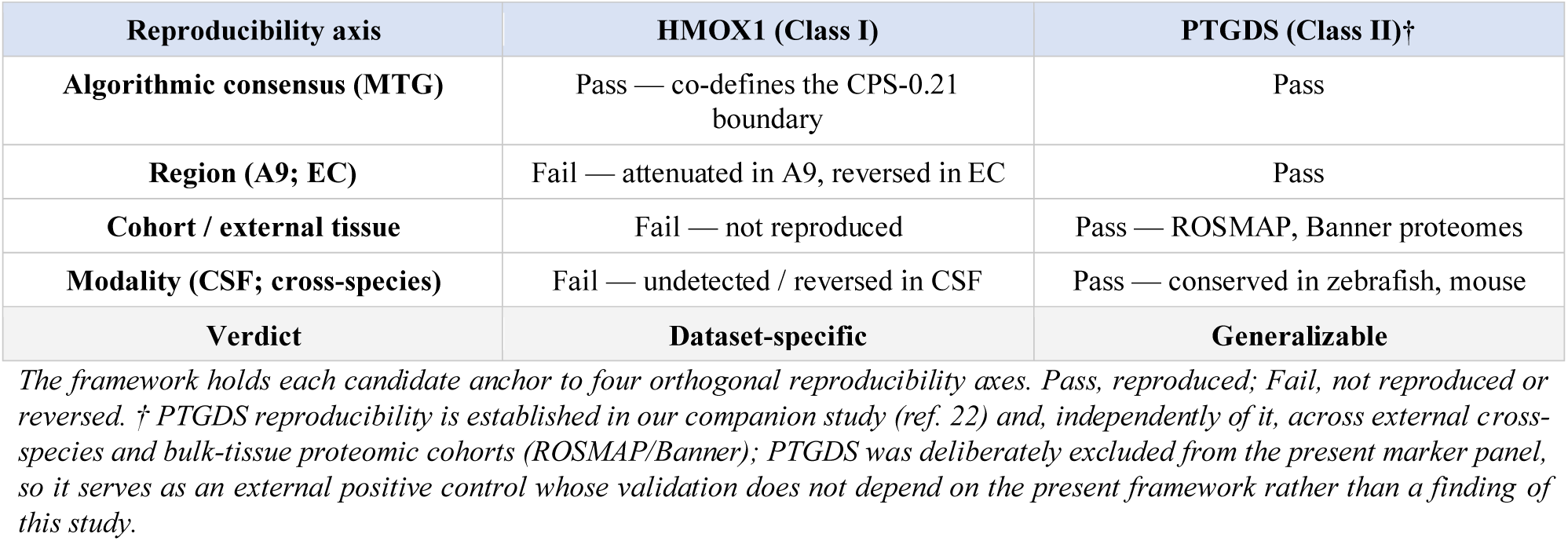
Marker validation across the four reproducibility axes.

**Table S7.** Donor-level adjudication of fourteen candidate AD markers in SEA-AD MTG.

| Marker | Cell type | CSF | Brain (lit.) | SEA-AD $\rho$ | SEA-AD trend | Reproduces? |
| --- | --- | --- | --- | --- | --- | --- |
| NPTX2 | Neuron | ↓ | ↓ | -0.58 | ↓ | ✓ |
| VGF | Neuron | ↓ | ↓ | -0.38 | ↓ | ✓ |
| SCG2 | Neuron | ↑/ns | ↓ | -0.24 | ↓ | ✓ |
| NPTX1 | Neuron | ↓ | flat* | +0.44 | ↑ | inversion (mRNA↑/protein↓) |
| NPTXR | Neuron | ↓ | flat* | -0.23 | ↓ | ✗ |
| VAMP2 | Neuron | ↑ | ↓ | +0.22 | ↑ | ✗ |
| NRGN | Neuron | ↑ | ↓ | +0.16 | flat | ✗ |
| SNAP25 | Neuron | ↑ | ↓ | -0.15 | flat | ✗ |
| CNTN2 | Neuron | ↓ | ↓ | +0.16 | flat | ✗ |
| GFAP | Astrocyte | ↑ | ↑ | +0.10 | flat | ✗ |
| CHI3L1 | Astrocyte | ↑ | ↑ | +0.08 | flat | ✗ |
| TREM2 | Microglia | ↑ | ↑ | -0.12 | flat | × |
| PTGDS | Astrocyte | — | ↑ | -0.31 | biphasic | biphasic (peak~0.47) |
| HMOX1 | Astrocyte | — | ↑ | -0.05 | flat | × |

For each marker the literature CSF and brain directions are compared with the SEA-AD donor-level transcriptional trend (Spearman $\rho$ of donor mean expression versus CPS, cell type as indicated). A marker counts as reproducing only when its SEA-AD direction matches the literature brain direction. Trends are donor-level monotonic Spearman $\rho$ versus CPS and are distinct from the boundary $\Delta z$ (a localized step at CPS ≈ 0.21); a marker such as GFAP can show a sharp boundary $\Delta z$ yet a weak donor-level monotonic trend. Only NPTX2, VGF and SCG2 reproduce; NPTX1 shows a compartment inversion (neuronal mRNA up while CSF/brain protein falls). Asterisk: NPTX1/NPTXR brain entries are the protein-level preserved prediction. Full statistics and code are in the reproducibility package (external\_marker\_panel\_audit.R, external\_marker\_audit.R).

